# Teaching a computer to assess hypnotic depth: A pilot study

**DOI:** 10.1101/2021.11.13.467562

**Authors:** Nikita V. Obukhov, Peter L.N.Naish, Irina E. Solnyshkina, Tatiana G. Siourdaki, Ilya A. Martynov

## Abstract

The therapeutic effects of hypnosis in some cases seem to be most marked when the patient has achieved sufficient hypnotic depth. It could be possible to monitor the deepening process using electrophysiological data to obtain information on depth changes throughout the session. However, although hypnosis is characterized by some common EEG patterns, significant differences between subjects are also observed. Therefore, an individualized approach is required to quantify the depth continuously during a session. To achieve this, we proposed the machine learning approach, using an EEG-based Brain-Computer interface, and tested it on video-EEG recordings of 8 outpatients. Based on the data from the first sessions, we trained the classification models to discriminate between conditions of wakefulness and deep hypnosis. Then, we applied them to subsequent sessions to predict the probability of deep hypnosis, i.e., to continuously measure depth level in real time. The models trained using frequency ranges of 1.5-14 and 4-15 Hz provided high accuracy. The applications and perspectives are discussed.

## Introduction

One of the differences between the procedures by which hypnosis is researched, compared with when it is applied in therapeutic settings, is whether or not hypnotizability is assessed. When researching the nature of hypnosis, it is common practice to include the use of a hypnotizability scale (Elkins, 2021). In contrast, it is very much the exception to find scales being used in clinical practice. There are many possible reasons for this, one of which could be the significant amount of time required to administer the assessment. This difficulty may now have been remedied, since Lush et al. (2021) have developed a scale which is delivered and assessed by computer. This would make it possible for a patient to be assessed without requiring lengthy input from a busy clinician.

Whether or not this facility would be of use to the clinician is debatable. It is tempting to extrapolate from the observation that hypnosis enhances therapeutic outcomes (Kirsch et al., 1995) to conclude that the benefits would be greatest in patients who were especially responsive to hypnotic techniques. However, evidence shows that the correlation between hypnotizability scores and clinical outcome is, at best, weak. For example, Abrahamsen and Naish (2021) found no correlation with level of analgesia achieved. These authors suggested an explanation, proposing that the raised level of anxiety, likely to be present in a therapeutic context, facilitated a greater hypnotic response. This is to be contrasted with the situation during hypnotizability testing, which is not an anxiety-provoking procedure. Whether or not this is truly the process which eliminates clear, hypnotizability-based inter-patient differences, it seems likely that intra-patient changes remain an important factor in therapeutic outcome. In other words, we propose that positive therapeutic effects are most marked when the patient has achieved an optimal state of hypnosis. This is certainly the tacit belief of most practitioners, who generally begin a treatment session with a hypnotic induction, then watch for signs that the patient has reached a sufficiently deep level of hypnosis before proceeding with specific treatment-related suggestions.

At this point we wish to make clear that we intend to take no theoretical position regarding the state-like aspects of hypnosis, nor whether the concept of depth, as applied to hypnosis, has any special meaning. We note merely that hypnotic procedures are associated with alterations in brain behavior (Tuominen et al., 2021) and that these changes appear to take time to become established (Kasos et al., 2018).

Since, as explained above, it has become possible to computerize the delivery of a hypnotic induction and hypnotizability test, it suggests that it may be possible to follow the automated induction with computerized monitoring of the deepening process and detection of the optimal stage for delivering suggestions. In theory, the entire treatment, therapy included, might be automated (and perhaps it will eventually be so, for certain kinds of treatment) but our interest is in the induction and depth assessment stages. To automate this would arguably be of far more value to a clinician than having computer-delivered hypnotizability tests. For one thing, it is common to give a patient practice, with several sessions of hypnosis, before the therapeutic procedure is attempted. Using hypnosis in childbirth would be an example of this, and busy medical staff could simply leave a woman with the computer, for a few occasions before the expected due date, gradually building and assessing her hypnotic skills. The automated depth assessment would also be valuable for a surgeon, who occasionally employed hypnosis for sedation and/or analgesia. With only occasional use of hypnosis, the surgeon is likely to be relatively inexperienced in looking for depth signs. We could imagine a case where a patient has already practiced with the “hypnosis equipment” and on the day of the operation uses it once again, in the operating theatre. When the surgeon enters, the patient is already approaching an ideal state, and it is necessary only to watch the computer monitor, to determine when it detects that the patient is ready.

Is such a scenario feasible? We argue that it might be if an appropriate means of detecting hypnotic depth were available. Naish (2010) showed that hypnosis was associated with a shift in hemispheric asymmetry, with an increase in right hemisphere processing and decrease in the left. This would be easy to monitor, but not by using temporal order judgments, as in the Naish study. Hemispheric shifts were also monitored in the Kasos et al. (2018) study, and in their case electrodermal measurements were employed; this would be an ideal approach in a clinical setting. However, it is unlikely that hemispheric changes can provide the basis of a universally satisfactory metric for assessing depth, because the asymmetry effect is very slight in people of low hypnotizability (Naish, 2010). Moreover, the shift can occur in the absence of hypnosis if a person becomes anxious (Shilton et al., 2019). It appears unlikely that an effective surrogate for neural changes will be found, so the only alternative is to monitor brain activity directly. Techniques such as MRI are not practical outside the laboratory, so EEG appears to be the only candidate. The difficulty here is that, although hypnosis is associated with changes in neural activity, the precise, fine-grained patterns observed differ from person to person. It would be possible for someone employing hypnotic techniques to learn the pattern typical of a specific patient, but to expect this for every patient would be contrary to the goal of automating these procedures. Moreover, simple recognition that the pattern was not optimal would not provide any quantitative information about the depth. The solution is to use machine learning. During the period when the patient was learning to become hypnotized, the machine would be learning to recognize their EEG patterns. This is the possibility that we explore in the current study.

Thus, **our research** aims to work out and test the system based on a machine learning approach to real-time control of the hypnosis depth level in patients, using an EEG-based Brain-Computer Interface (BCI).

## Materials and methods

The BCI used in this study was the passive type (Kroll et al., 2018; Roy et al., 2013; Zander et al., 2017), and supervised learning was employed (Kostenko et al., 2022; Liu & Wu, 2012). In this technique two or more mental states are used; we opted for two: waking and deeply hypnotized. The first session with each patient was the calibration (training) one when the machine learned to discriminate between two states. To provide this training, we had to define and label manually several epochs of an obtained EEG recording that corresponded to these two states.

Waking was relatively simple to define; hypnosis was rather more subjective. An experienced hypnotist was used, first to establish a reasonable level, then to deepen by getting the patient to visualize descending a series of steps, counting as they went. To select periods of deep hypnosis two reasonably objective measures were employed. The first picked times when the face muscles became “smooth”, the lowering of the mandible (sometimes accompanied by slight mouth opening) and a decrease in frequency of breathing were observed (Casiglia et al., 2018; Casiglia et al., 2019). The other was based on spontaneous post-hypnotic amnesia, which was detected retrospectively. After waking, the patient was questioned about the session and was judged to have been amnesic from the point where they last remembered walking down steps, to the point where they remembered walking up again. Both criteria had to be present to be judged deep.

Each session was video captured synchronously with EEG recording for better detection of the times when physical signs of deep hypnosis appeared. For the training, within each of the two types of periods in the calibration recording, we placed event labels corresponding to the period: event “W” (for the wakefulness) and event “D” (for the deep state), so that they could be used to specify of the order of 20-25 short trial epochs for each state. We attached the “D” marks to the moments of the visual signs of sufficient depth within the periods of reported amnesia. The “W” labels were placed arbitrarily along the phases of baseline EEG registration.

Therefore, using two sets of epochs, the system learned to recognize (“predict”) the waking and deeply hypnotized states in the subsequent sessions online (i.e., in real time) with some interdependent probability. This could be interpreted as measuring the hypnotic depth - the higher the probability of the deepest stage, the deeper the hypnosis.

The machine learning system comprised equipment for synchronized recording of EEG, video, and audio, together with various software packages. The hardware and electrode placement are detailed in “Supplementary Materials A, part 1”.

EEG spectral analysis was performed in WinEEG software, version 2.130.101 (WinEEG Research Software, n.d.), and marking the calibration EEG recording for classifier training was carried out in EEGLab 2019.0 (Delorme & Makeig, 2014). The construction of designs for the offline training, online application of trained prediction models, and classification accuracy testing was carried out in OpenVibe 2.2.0 (Renard et al., 2010).

### Participants

Nine patients (3 men and 6 women, mean age 38.33 ± 10.61 years) underwent up to 7 hypnosis sessions with synchronized EEG and video recording. The inclusion criteria were the age between 18 to 65 years, the presence of a diagnosis of anxiety or comorbid anxiety, depressive or somatoform disorders (to homogenize the sample) that met the corresponding ICD-10 criteria, and consent to participate in the study. All patients filled out an informed consent form approved by the Association of Experts in the Field of Clinical Hypnosis (Saint Petersburg) Ethics Committee. The exclusion criteria were severe cognitive or somatic impairment, anamnesis of epilepsy, psychotic episodes, and the absence of posthypnotic amnesia periods in the first session. The summary of participants is presented in Table 1.

**Table 1.** Participants’ data

| **Patient's Code** | **Gender** | **Age, years** | **Diagnosis, ICD-10** | **Number of sessions** | **Medications, mg per day** |
| --- | --- | --- | --- | --- | --- |
| A | M | 34 | F41.1<br>Generalized anxiety disorder+ F45.8<br>Other somatoform disorder | 7 | Tofisopam, 100 |
| E | F | 58 | F40.0<br>Agoraphobia | 6 | Escitalopram, 10 |
| G | F | 45 | F41.1<br>Generalized anxiety disorder | 4 | Sertraline, 50 |
| S | F | 49 | F40.01<br>Agoraphobia with panic disorder | 3 | Escitalopram, 10 |
| O | F | 33 | F32.1<br>Moderate depressive episode+ F41.1<br>Generalized anxiety disorder | 2 | Fluoxetine, 40 |
| N | M | 35 | F41.1<br>Generalized anxiety disorder | 2 | Fluoxetine, 20 |
| V | M | 35 | F41.1<br>Generalized anxiety disorder | 2 | - |
| C | F | 22 | F32.1<br>Moderate depressive episode+ F41.1<br>Generalized anxiety disorder | 2 | Sertraline, 100 |
| T | F | 34 | F41.1<br>Generalized anxiety disorder | 1 | - |

### EEG files

EEG files were stored in .gdf format. The initial number of EEG recordings was 29. Recordings of poor quality (a large number of artifacts, high electrode impedance) were excluded from the analysis.

### Session procedure

The electrodes were placed to register the patient’s EEG, and the equipment was installed for video and audio recording to be synchronized with EEG so that the video captured the head, trunk, and upper limbs. Then, a baseline EEG was recorded with the eyes closed for 3-5 minutes.

Then followed the hypnosis and deepening, as detailed above. Suggestions were additionally given to maintain or deepen the hypnosis, together with rapport checking, by giving the suggestion, “If you hear me, let your unconscious mind move the thumb of your right hand.” Further patient-specific suggestions were then given, then finally the patient was counted back.

### Analysis of the obtained EEG recordings synchronized with the video

The collected data were processed in four different ways. The first did not use machine learning, but underpinned it, by ensuring that the electrophysiological distinction between the waking and deep hypnosis remained constant in each patient throughout the treatment.

For this type of analysis, artifacts in the EEG recordings were deleted with the Infomax ICA decomposition (Langlois et al., 2010). Using WinEEG we then compared the averaged power spectra of the identified deep sections and the waking sections (approx. 5-10 min each) in the Delta (1.5-4 Hz), Theta (4-8 Hz), Alpha (8-12 Hz), Sensory-motor or Low beta (12-15 Hz), Beta1 (15-18 Hz), Beta2 (18-25 Hz) and Gamma (25-45 Hz) frequency bands. This provided general information about which rhythms in which brain areas changed after achieving deep hypnosis in each session.

The second type of data processing was the machine learning procedure, based upon the “W” and “D” labeling described previously. As a result, we trained prediction models to be used in the subsequent sessions for the online classification. To determine what frequency band could give us the trained models with the most predictive power, we used a wide frequency band (1.5 - 45 Hz) and also three narrower bands: 1.5-8 Hz, 1.5-14 Hz, and 4-15 Hz, since lower frequencies activity has been shown to be most during hypnosis (Cordi et al., 2014; De Benedittis, 2021; Jensen et al., 2018; Wolf et al., 2022). We improved the classification accuracy by spatial filtering of signals with the Common Spatial Pattern (CSP) method (Aydemir, 2016; Bird et al., 2019; Blankertz et al., 2008; Ramoser et al., 2001), then we assessed the accuracy of 4 types (according to the selected bands) of models by the 10-fold cross-validation test for each session. “Supplementary materials A, Part 2” contain a detailed technical description of this type of analysis.

The third process utilized trained models to make a real-time assessment of the second and subsequent sessions. This produced a Probability Value parameter, varying between 0 and 1, depending on the similarity to the learned pattern of deep hypnosis. This parameter was displayed in the form of a curve on the Continuous Oscilloscope and informed the hypnotist of the likely depth of the hypnosis. We called this curve the Predictive curve. For a detailed description of this step of analysis see “Supplementary materials A, Part 3”.

Finally, we checked the accuracy of the four types of prediction models based on the current “W” and “D” labeling of each second and subsequent session. These labels were placed by someone not privy to the predictive outcomes of the sessions. Then, we calculated the accuracy as a percentage of coincidence of the states predicted by the model for different epochs with actual (based on the labels) states related to these epochs (Kohavi & Provost, 1998).

The training on the data of these sessions resulted in obtaining new models. They were not used further online, but we applied them to classify the same data on which these models had been trained. This way after each second and subsequent session we built a curve that reflected their dynamics with maximal accuracy. We called it the Native curve. Then, the Native and Predictive curves of the same sessions were visually compared to additionally assess how accurately the Predictive curve describes the real dynamics. This indicated the ongoing stability of the technique. For technical details of this part of the analysis see “Supplementary materials A, Part 4”.

## Results

All the recordings were free of oculographic artifacts due to the fact that the patients already had their eyes closed at the very beginning of EEG registration. Patient T was excluded from the study because there were no periods of amnesia as the session result. Also, one of the recordings of Patient E (session # 5) was excluded from the analysis due to the increasing myographic artifacts. Therefore, the total number of EEGs from the remaining 8 participants was 27.

### Results of the assessment of deep hypnosis patterns in different patients and their constancy throughout the treatment

These results are presented as topographic maps showing the degree of differences in the averaged power spectra between the hypnotic period and the awake phase of EEG recording (“deep hypnosis” minus “wakefulness” spectra). Thus, the maps demonstrate the changes in the power of different rhythms for different localisations after achieving deep hypnosis. Figure 1 shows examples of such maps for three patients at their three different sessions. “Supplementary materials B” contain all of these maps for all 27 included sessions of all patients.

**Figure 1.**
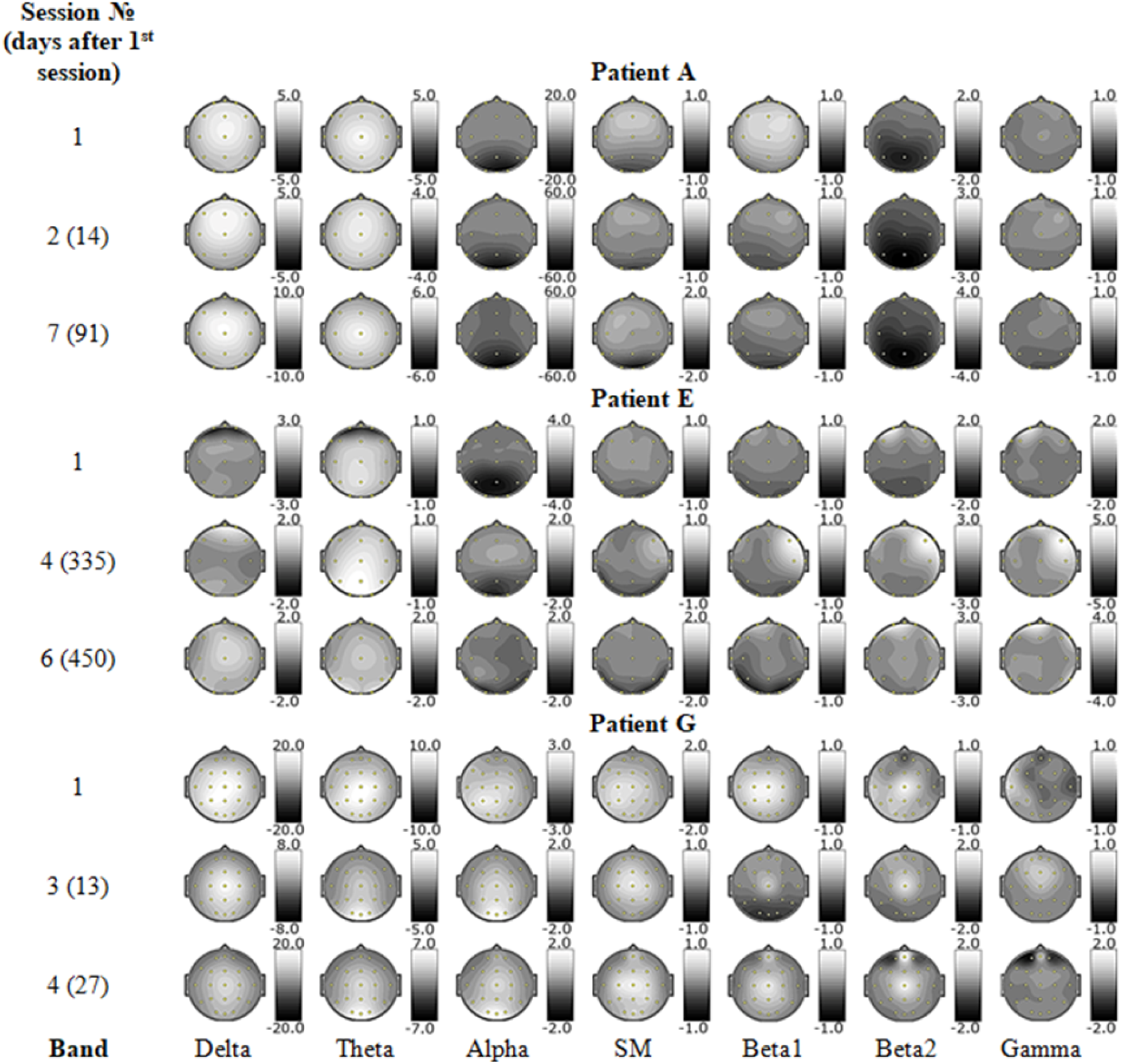
An example of topographic maps displaying the changes in the power of different rhythms for different localisations after achieving deep hypnosis for three patients (sessions for displaying are randomly selected). Changes in the power of rhythms are displayed in color according to the graduation of a nearby color scale. The color scale is in μV^2^

### Results of the 10-fold cross-validation test of wakefulness and deep hypnosis discrimination accuracy

Table 2 shows the results of this test for four types of classification models according to the band used to train the model. It is common to display the results of such analysis in the form of the mean value of the classification accuracy among the ten partitions of the marked file, and the corresponding standard deviation. Despite its necessity, this test cannot be comprehensive, because it assumes accuracy checking on the same data which are the training ones, hence it was also necessary to test the prediction model accuracy on new data, i.e., data from the second and subsequent sessions.

**Table 2.** Accuracy of classification of the wakefulness state and deep hypnosis for four types of prediction models according to the band used to train the model: The 10-fold cross-validation test results

| Patient | Session | Accuracy for the corresponding band, M±SD, % | | | |
| --- | --- | --- | --- | --- | --- |
|  |  | 1.5-45 Hz | 1.5-8 Hz | 1.5-14 Hz | 4-15 Hz |
| A | 1 | 96±8 | 86±18 | 98±6 | 94±9.17 |
| | 2 | 93.5±10.01 | 95.5±9.07 | 86.5±14.33 | 72.5±26.48 |
| | 3 | 88±18.33 | 72±31.24 | 86±25.38 | 86±25.38 |
| | 4 | 95±15 | 83±15.84 | 95±10 | 95±10 |
| | 5 | 94±9.17 | 89.67±17.16 | 92.33±13 | 96.33±7.37 |
| | 6 | 98±6 | 87±15.52 | 92±13.27 | 96±8 |
| | 7 | 84.17±21.87 | 93.33±13.33 | 83.33±22.36 | 86.67±16.33 |
| | M±SD | 92.67±4.86 | 86.64±7.75 | 90.45±5.32 | 89.5±8.65 |
| E | 1 | 97.5±7.5 | 82±15.68 | 88.5±16.13 | 95±10 |
| | 2 | 100±0 | 90.5±11.72 | 93±15.52 | 86.5±19.11 |
| | 3 | 100±0 | 91±11.14 | 100±0 | 100±0 |
| | 4 | 95±10 | 93±15.52 | 91±15.78 | 92±24 |
| | 6* | 100±0 | 96±8 | 94±18 | 96±8 |
| | M±SD | 98.5±2.24 | 90.5±5.22 | 93.3±4.3 | 93.9±5.03 |
| G | 1 | 94.33±8.7 | 96±8 | 98.33±5 | 96±12 |
| | 2 | 98.33±5 | 92.33±18.14 | 94.67±11.08 | 93±15.52 |
| | 3 | 100±0 | 100±0 | 100±0 | 96.33±7.37 |
| | 4 | 98±6 | 98±6 | 98±6 | 98±6 |
| | M±SD | 97.67±2.39 | 96.58±3.27 | 97.75±2.23 | 95.83±2.08 |
| S | 1 | 100±0 | 100±0 | 97.5±7.5 | 95.5±9.07 |
| | 2 | 97.5±7.5 | 100±0 | 100±0 | 100±0 |
| | 3 | 97.5±7.5 | 97.5±7.5 | 97.5±7.5 | 97.5±7.5 |
| | M±SD | 98.33±1.44 | 99.17±1.44 | 98.33±1.44 | 97.67±2.25 |
| O | 1 | 100±0 | 100±0 | 97.5±7.5 | 97.5±7.5 |
| | 2 | 100±0 | 100±0 | 97.5±7.5 | 87.5±16.78 |
| | M±SD | 100±0 | 100±0 | 97.5±0 | 92.5±7.07 |
| N | 1 | 96±8 | 98±6 | 94±18 | 98±6 |
| | 2 | 100±0 | 100±0 | 100±0 | 100±0 |
| | M±SD | 98±2.83 | 99±1.41 | 97±4.24 | 99±1.41 |
| V | 1 | 91±15.78 | 93.5±10.01 | 98±6 | 95.5±9.07 |
| | 2 | 100±0 | 98±6 | 100±0 | 100±0 |
| | M±SD | 95.5±6.36 | 95.75±3.18 | 99±1.41 | 97.75±3.18 |
| C | 1 | 100±0 | 95±15 | 81.67±22.91 | 96.67±10 |
| | 2 | 100±0 | 77.5±29.36 | 93.33±20 | 93.33±20 |
| | M±SD | 100±0 | 86.25±12.37 | 87.5±8.24 | 95±2.36 |

\* session 5 has been excluded from the analysis

### Results of visual (qualitative) testing of the proposed methodology in real time (in the second and subsequent sessions)

As an example of how the prediction model works in real time, Figure 2 shows a screenshot of the graph (the Predictive curve) reflecting changes in time of the probability that the person is in a deep hypnotic state (effectively, the hypnotic depth). It was taken at the end of the online session with Patient A on the seventh visit, the last and shortest for him. “Supplementary materials C” contain the screenshots of the Predictive curves (the upper ones) derived for each patient of the study from the second and the following sessions using a suitable classification model.

**Figure 2.**
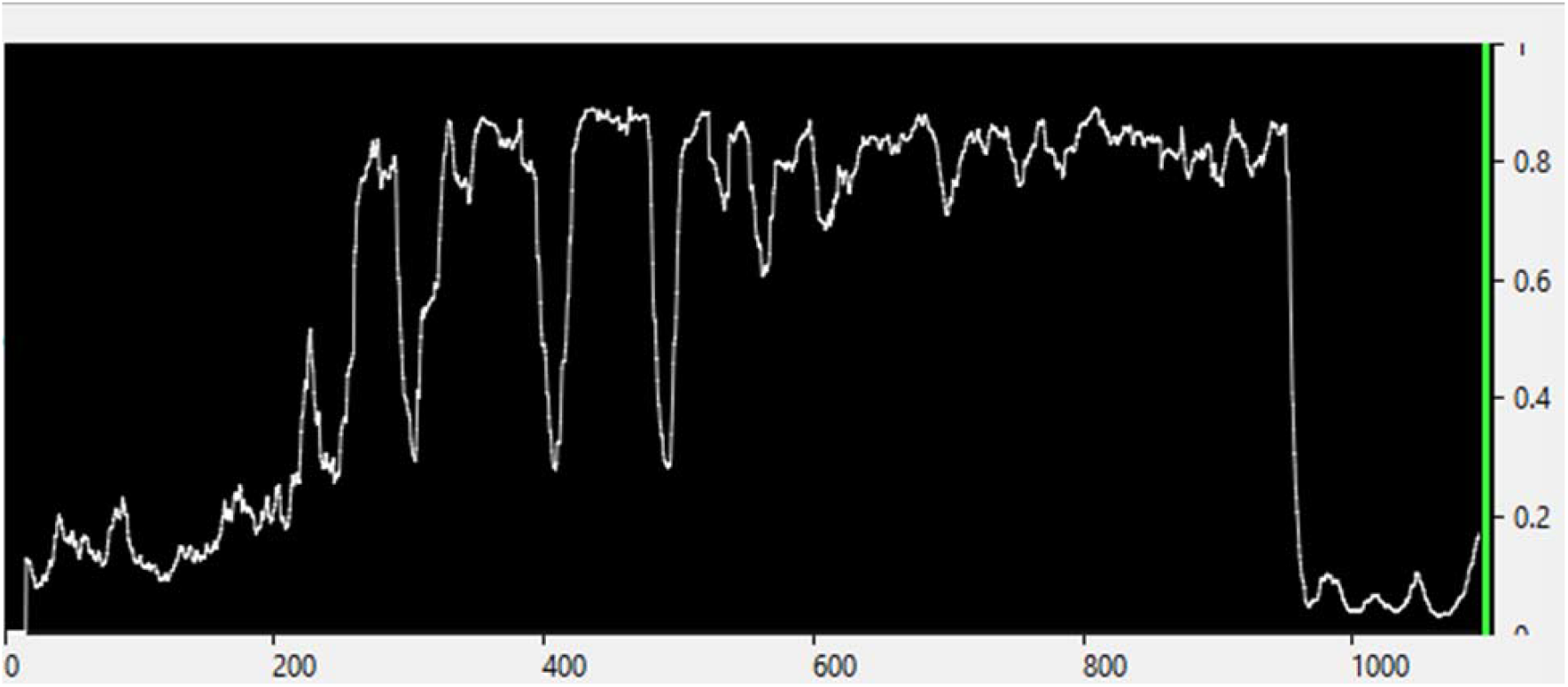
An example of the Predictive curve. Screenshot of the curve at the end of the online session with Patient A on the seventh visit. In this example, the classification model was trained using the 1.5-14 Hz range. The curve was smoothed by the Moving epoch average (Immediate) function. The number of 4-s epochs with an overlap of 0.5 s used for averaging was 50

Qualitatively comparing the curves data with the patients’ physical signs of being awake and being deep, we found an approximate coincidence of the time when the curve was consistently around 0.7 or above (on average, depending on smoothing features), with a completely amnesic period and the presence of deep hypnosis peripheral signs. The sections of the high-amplitude wave-like motion of the curve, in general, were associated with a report of alternating awareness and partial amnesia for this period. Generally, the periods of the more or less constant curve position below approximately 0.6 were characterized by the patients as a deep bodily relaxation with an inability to open the eyes and move the limbs, total awareness, and no amnesia. And the closer to 0, the more signs of total awake state the patients reported.

### Results of quantitative evaluation of the classification accuracy based on the data of the second and subsequent sessions and visual comparison of the Predictive and Native curves

The results of this type of accuracy estimation are shown in Table 3. As had been expected, the accuracy is lower than that evaluated by the 10-fold cross-validation test, and this reflects the natural variability of the electrophysiological characteristics of the different sessions. Nevertheless, the classification models from Patients G, N, and V for all four frequency ranges gave very high average accuracy, in some cases reaching values above 95%. In Patients A and O, only one of the four bands (1.5-8 Hz for Patient A and 1.5-45 Hz for Patient O) gave unsatisfactory results, while all the others had high predictive power. Two bands gave good results in Patients S (1.5-8 and 1.5-14 Hz) and C (1.5-45 and 4-15 Hz). The greatest difficulties occurred only with Patient E, where the classification accuracy was quite high only in the 4-15 Hz band. Figure 3 represents accuracy averaged over all sessions (except the calibration ones) with the standard deviation (SD) for four bands. As seen, the highest accuracy has been obtained using 1.5-14 and 4-15 Hz. The lowest SD for 4-15 Hz shows that this band provided the most stable accuracy across different sessions.

**Table 3.** Accuracy of classification of the wakefulness state and deep hypnosis based on the data of the second (and subsequent) sessions for four types of prediction models according to the band used to train the model

| Patients | Sessions | Accuracy for the corresponding band, % |  |  |  |
| --- | --- | --- | --- | --- | --- |
|  |  | 1.5-45 Hz | 1.5-8 Hz | 1.5-14 Hz | 4-15 Hz |
| A | 2 | 74.31 | 52.08 | 74.65 | 66.32 |
|  | 3 | 80.0 | 75.34 | 91.51 | 89.86 |
|  | 4 | 99.48 | 64.83 | 87.66 | 88.98 |
|  | 5 | 69.09 | 66.94 | 94.62 | 91.94 |
|  | 6 | 90.85 | 69.82 | 90.24 | 87.8 |
|  | 7 | 79.73 | 66.32 | 80.76 | 73.54 |
|  | M±SD | 82.24±11.12 | 65.89±7.72 | 86.57±7.48 | 83.07±10.52 |
| E | 2 | 54.75 (98.42)* | 38.29 | 70.89 | 80.38 |
|  | 3 | 54.18 (72.76)** | 65.63 | 53.56 | 83.9 |
|  | 4 | 71.82 | 69.7 | 81.52 | 84.55 |
|  | 6*** | 70.06 | 44.19 | 59.01 | 84.01 |
|  | M±SD | 62.7±9.54 | 54.45±15.53 | 66.24±12.49 | 83.21±1.9 |
| G | 2 | 87.10 | 91.13 | 91.13 | 73.92 |
|  | 3 | 88.98 | 97.31 | 90.05 | 81.72 |
|  | 4 | 75.78 | 94.3 | 89.46 | 87.75 |
|  | M±SD | 83.95±7.14 | 94.25±3.09 | 90.21±0.85 | 81.13±6.93 |
| S | 2 | 53.03 | 93.64 | 95.76 | 53.64 |
|  | 3 | 94.3 | 87.03 | 78.16 | 75.95 |
|  | M±SD | 73.67±29.18 | 90.34±4.67 | 86.96±12.45 | 64.8±15.78 |
| O | 2 | 61.46 | 89.58 | 89.24 | 88.89 |
| N | 2 | 97.09 | 93.6 | 96.8 | 96.51 |
| V | 2 | 95.06 | 90.41 | 88.08 | 94.48 |
| C | 2 | 71.92 | 61.15 | 66.15 | 82.31 |

\* value after a drift correction with subtraction of 1.5 from the original feature vector (for details see "Supplementary materials D")
\*\* value after a drift correction with subtraction of 2.5 from the original feature vector (for details see "Supplementary materials D")
\*\*\*session 5 has been excluded from the analysis

**Figure 3.**
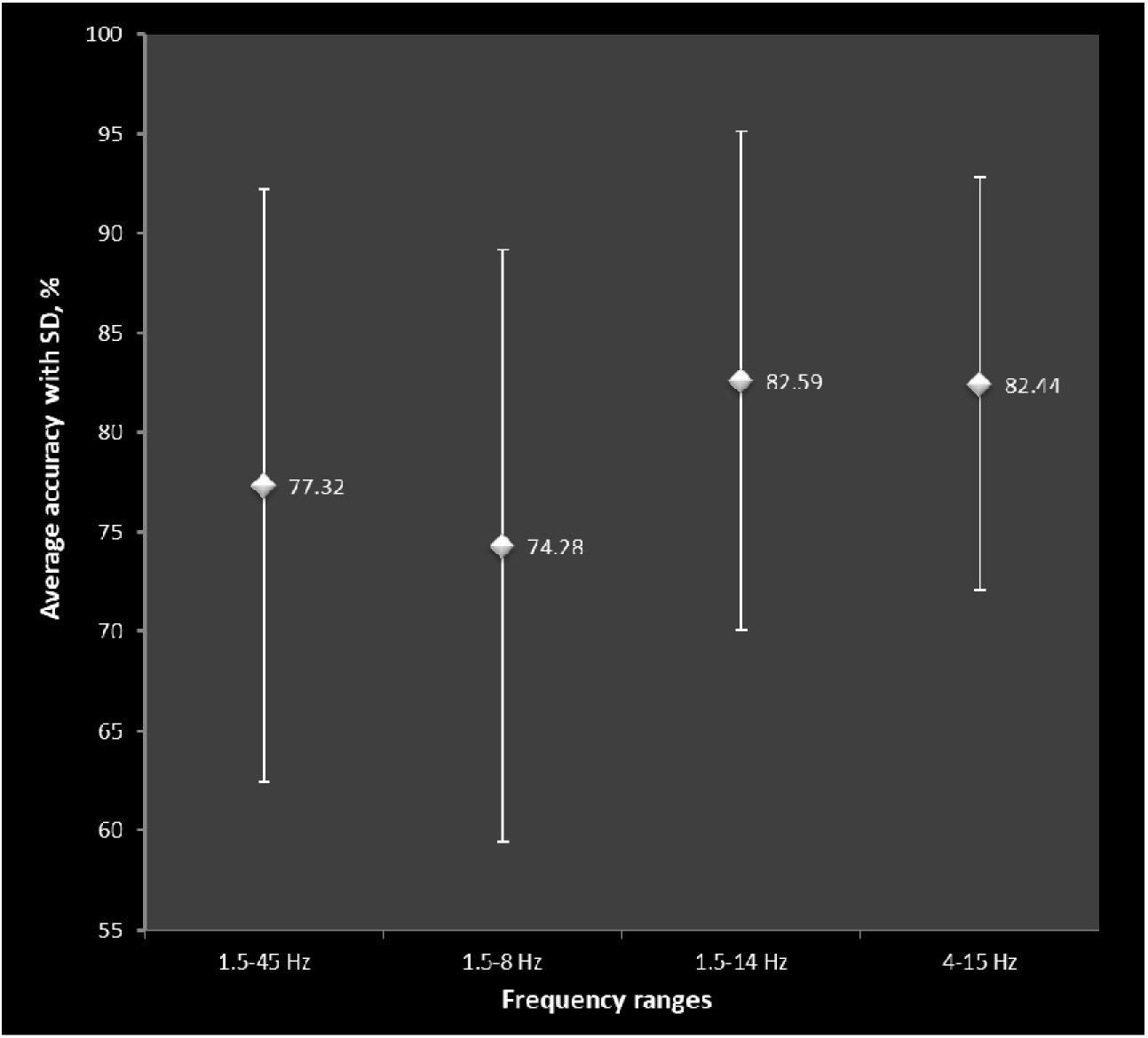
Accuracy of classification of the wakefulness state and deep hypnosis based on the data of the second and subsequent sessions (averaged over all these sessions) for four bands used to train the model. Vertical lines reflect the SD

As an example of the results for visual comparison of the Native and Predictive curves of hypnosis depth dynamics, Figure 4 shows the Native curve of Patient A’s seventh session. By comparing the graphs in Figure 2 (the Predictive curve) and Figure 4 (the Native curve) we can see that the configurations of the curves largely coincide, giving us additional confirmation that the prediction model can reflect the real picture more or less accurately. “Supplementary materials C” contain the pairs of the Predictive and Native curves for each patient of the study for the second and the following sessions which demonstrates the noticeable coincidences between the curves within each pair, if a model of high classification accuracy was used to get the curves.

**Figure 4.**
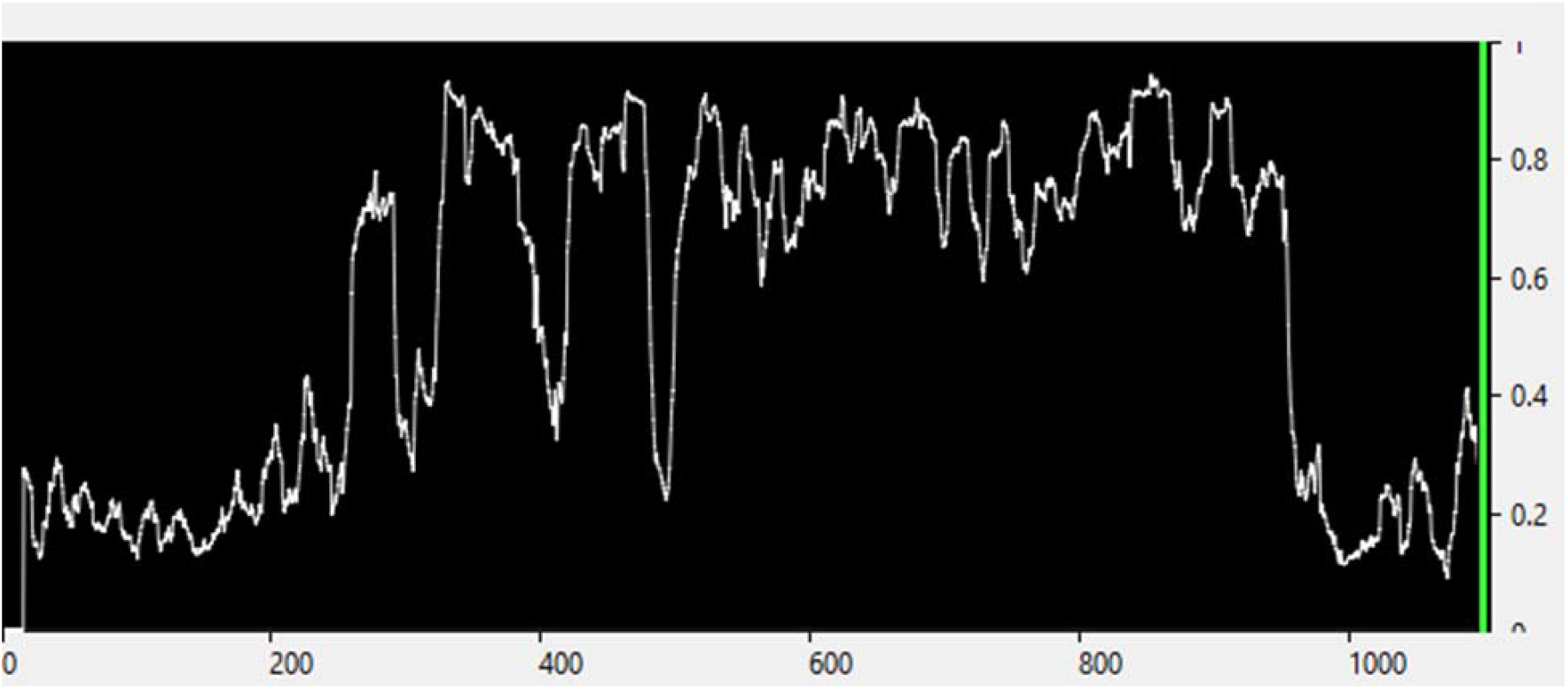
An example of the Native curve. Screenshot of the Native curve of the seventh session with Patient A. The band and the smoothing features are the same as in the Predictive curve

## Discussion

The results of the first step of our analysis, which are presented in Figure 1 and “Supplementary materials B”, provide approximate information about which rhythms in which brain areas changed after achieving deep hypnosis in a particular session. For example, as we can see from the topographic maps of Patient A, his transition to a deep hypnotic state is associated with a very large decrease in the alpha range activity, some decrease in the beta-2 rhythm in the occipital regions, a noticeable increase in theta, and even slower activity in the middle and mid-frontal regions, some increase of the sensorimotor and beta-1 rhythm in the frontal region. This combination tends to be observed in each of his sessions. The maps of the other patients demonstrate some similar changes: the slow-wave activity increase in different areas of their brain, which corresponds to the literature on the electrophysiological correlates of hypnosis (Cordi et al., 2014; De Benedittis, 2021; Deivanayagi et al., 2007; De Pascalis et al., 1998; De Pascalis & Santarcangelo, 2020; Freeman et al., 2000; Jamieson & Burgess, 2014; Jensen et al., 2015, 2018; Kropotov, 2009; Paoletti et al., 2020; Sabourin et al., 1990; Vaitl et al., 2005; Wolf et al., 2022). Nevertheless, they also show quite clear differences between patients. For example, Patient G, unlike all the other participants of the study, did not demonstrate a decrease in the alpha rhythm in the occipital region; on the contrary, she showed an increase alongside an increase in the power of slower rhythms.

Also, we can see from the maps that the patterns of deep hypnosis of a particular patient show noticeable stability over time and tend to be reproduced from session to session. The example of Patient E demonstrates that this reproducibility can be observed even if more than a year passes between the measurements. Thus, based on the analysis of the topographic maps of all 27 hypnosis sessions, we can make general preliminary conclusions that: a) the patterns of deep hypnosis brain activity are reproduced from session to session in a particular patient, which suggests that b) in general, using the EEG recording of any patient’s session as a calibration file, we can expect the previously trained system to detect the desired states in subsequent sessions. We can also conclude that, although the deep state patterns may have certain similarities among patients, c) there are noticeable differences between the subjects, which tells us that d) it is impossible to use any general approach to evaluate and “measure” hypnotic depth for all people, and an individualized assessment is required. All of these points mean that the machine learning approach can potentially apply to hypnosis.

As seen from Table 2, the classification accuracy obtained by the 10-fold cross-validation tests in all patients and all the tested frequency ranges is quite high, in the vast majority of cases, exceeding 85%. This indicates that the trained system can discriminate well between two states within each EEG recording using the chosen algorithms. It also underpinned the results of the previous type of analysis, showing that patterns of deep hypnosis are quite different from waking ones. For some sessions, the accuracy turned out to be absolute. Possibly, this was due to the tendency of the OpenVibe software to give optimistic calculation results, and the manufacturer warns about this (Renard et al., 2010). It is also probable to be due to the over-fitting phenomenon (Santos et al., 2018), which was taken into account by us. Consequently, despite the cross-validation test being quite a reliable method, these results were important but not comprehensive. Therefore, we performed other estimations of our system feasibility.

The moment by moment probability values of deep hypnosis were provided as a form of feedback for the hypnotherapist, and were displayed in real time during second and subsequent sessions in the form of a Predictive curve. We were able to control its motion, and therefore not only predict the deepest (amnesic) periods but also evaluate how hypnosis was developing between the total wakefulness and deepest stages. It is difficult to assess the fine-grained depth dynamics by the physical signs, therefore such instrumental control could be effective. As a result of qualitative analysis based on post-session patients’ reports, we found that the curve generally can reflect the phenomenology of the session.

To demonstrate this in detail we can use the example from Figure 2. Here we can highlight several quite distinct periods in the curve of Patient A. For about the first 200 seconds, the probability values of the deep state are at a low level, which corresponded to calm wakefulness with eyes closed. Then suggestions for hypnosis induction followed. Since it was the seventh session with this patient, and he had already developed the skill to quickly achieve a deep state, the curve rises quite abruptly, but for some time (from about 200^th^ to 500^th^ seconds) it is undulating. This was expressed in the patient’s further report that he periodically “plunged deeper”, and “switched off”, but briefly “surfaced” from time to time and that he remembered this session part in fragments (the partial amnesia). This part of the session probably could be called the period of development (formation) of deep hypnosis. From about the 500^th^ second, the curve values stabilize around 0.8, keeping only minor fluctuations, and the patient was completely amnesic about this part of the session. Concise therapeutic suggestions had been made and, since about the 900^th^ second, the awakening phase had started by the method of reverse counting from “5” to “1” with full awakening on the count of “1” (without opening the eyes to avoid oculographic artifacts). This phase lasted for about 1 minute. However, as can be seen from the curve, up to the moment when the hypnotherapist pronounced the number “1” (959 s), the patient had been in a deep state. Immediately after that, there was a sharp exit, followed by some time of baseline EEG registration with the eyes closed (from 960 to 1105 seconds).

The Predictive curves of all the rest of the participants and sessions (“Supplementary materials C”, the upper screenshots) describe generally the peculiarities of each session by the same principle. A more or less accurate reflection of the session phenomenology in the Predictive curve configuration, based on post-session patients’ reports, could indirectly indicate that our approach may have high predictive power. This judgment can be underpinned by comparing the graphs of the Predictive and the Native curves of each online session of the patients (Figures 2 and 4, and “Supplementary materials C”). We can see that their configurations related to the particular session coincide, sometimes greatly, sometimes with acceptable variations, if the model of high classification accuracy was used to get the curves. For example, Figure 4 (the Native) shows the periods of hypnotic depth dynamics that are the same as those we were able to observe on the Predictive curve.

Table 3 shows that the classification models trained in the first sessions, in general, accurately predicted the actual states of the patients in the next sessions. The classification models from Patients G, N, and V gave very high average accuracy for all four frequency ranges. In Patients A and O, three bands had high average predictive power. Two bands gave good results in Patients S and C. Patient E was an individual case of the study. The average classification accuracy was quite high for her only in the band from 4 to15 Hz. Probably, it was caused by some experimental issues, unique to her, and many EEG recording artifacts, e.g., the phenomenon of drift (trend) (Driel, 2021). For a detailed analysis of these case-related problems, in this and other cases of the study, as well as our conclusions on how to avoid them further, see “Supplementary materials D”.

Table 3 also shows that each patient had their own “preferred” frequency band where the classification model was the most accurate. For example, for Patient A it is 1.5-14 Hz, for Patients G and S it is 1.5-8 Hz, etc., which reflects the aforementioned individuality of the hypnotic state patterns. The calculation of the classification accuracy for each band by averaging the data over all sessions (except calibration ones) shows that the range from 1.5 to 14 Hz had the highest and quite stable accuracy. The 4-15 Hz band gave similar results with greater stability (see Figure 3).

Thus, the system designed in this study could be used to monitor the dynamics of the depth of hypnosis in a more refined and detailed way. The motion of the Predictive curve, its tendency to decrease or increase could give a lot of additional information. This raises the question of why we need this information and how we can use it.

Answering this question, first, we could say that, the system may be a prototype of a tool for quantifying hypnosis depth, “measuring” it, and analyzing its dynamics both during a session and after it. We can see more explicitly that depth can have some microstructure over the session duration, where some phases could be recognized: the period of dipping into hypnosis, the periods of forming, stabilization, and exit. It has some peculiarities in a particular case, e.g., a tendency to undulate during the forming period, a certain speed of deepening or awakening, etc.

Secondly, this system could contribute to the optimization of clinical work, which is expressed in various aspects. The central aspect is that the ability to control the level of depth of hypnosis more precisely allows us to influence its degree or to use knowledge about the moments of the maximal depth in practice. In the Introduction, we tried to show that deeper hypnosis may have several therapeutic benefits in particular cases. In addition, hypothetically it can be useful to reach the phenomenon of posthypnotic amnesia for the received suggestions to minimize the intervention of consciousness, critical attitude, and resistance to them.

Watching the Predictive curve can allow us to detect the correlation between the verbal or nonverbal therapist’s behavior and the reactions of the patient’s brain, which can help the therapist to flexibly respond according to the individual needs of the patient by selecting words, intonation, the pace of speech and other parameters of the session management. Additionally, we could probably immediately detect the initial signs of depth disturbance and restore it with the appropriate suggestion, as well as use ratification, when the curve movement shows the desired trend. Furthermore, there may be a hypothetical relation between a greater duration of deep periods in some particular session and a greater anxiety (or pain, etc.) reduction as a result of it (which certainly needs to be tested).

An important effect of using our system could be an increase in the competence of the specialist, which is expressed in the training of understanding (“feeling”) the hypnosis dynamics and strengthening the ability to more skillfully monitor the external signs that correlate with the changes in brain activity. This system can also be the basis for creating neurofeedback protocols for teaching patients to go into hypnosis on their own, where the feedback parameter will be the data of the Predictive curve presented in an auditory or tactile form.

## Conclusion

This study showed that the electrophysiological patterns of the deep hypnotic state are stable from session to session in a patient, even over long intervals. They reveal several similarities among patients, but at the same time are characterized by significant individuality.

It is possible to create a system based on the principle of passive BCI with supervised learning. A classification model trained on the data from a session can allow us to accurately predict the state of deep hypnosis in subsequent sessions, providing an individualized parameter that quantitatively describes the level of depth in real time.

Classification models trained using frequency ranges of 1.5-14 and 4-15 Hz can provide high accuracy. With this system, the hypnotherapist could monitor the session dynamics more precisely and use the data obtained to optimize therapy as well as theoretically study the tendencies of the depth to change depending on different circumstances. Furthermore, it could potentially be implemented as an individualized neurofeedback tool for self-hypnosis.

## Limitations

The limitations are the heterogeneous number of experimental sessions and the artifacts in some EEG recordings. Although, as shown, the problem of artifacts could be solved by careful frequency band selection for the classification model, further development of algorithms for eliminating artifacts in real time is desirable.

Our study involved only those patients who were able to achieve spontaneous posthypnotic amnesia. However, this phenomenon is just a particular case of some target conditions. Therefore, while training the classification model, a specialist can assign any state or phenomenon that they want to predict in the next sessions as a target.

Further research is necessary to test other feature-extraction approaches and alternative methods of classifying states (e.g., Support Vector Machine, Multilayer Perceptron, or ensembles of classifiers) (Bird et al., 2018; Bird et al., 2019).

Future development of our study also includes the analysis of the correlation of electrophysiological parameters with the hypnosis session clinical effects, e.g., anxiety reduction level as a result of the session. Moreover, we plan to work out self-hypnosis personalized protocols based on feedback using machine learning.

## Acknowledgments

We thank Gelera U. Utemisheva, Svetlana V. Khetrik, Valentin A. Kuznetsov, and Michael P. for their help with organizational issues.

## Data availability statement

The authors confirm that the data supporting the findings of this study are available within the article and its supplementary materials.

