## Supplementary materials A for "Teaching a computer to assess hypnotic depth: A pilot study"

**Part 1**

For EEG recording, we used a Mitsar-EEG-SmartBCI 21-channel neurointerface (CE medically certified (MDD 93/42 / EEC), the hardware sampling rate was 2000 Hz filtered down to 250 Hz, the frequency range was DC(0)-70 Hz) in combination with an elastic textile cap with fixing rings for point Ag/AgCl sintered electrodes located according to the international system 10-20 in positions: Fp1, Fpz, Fp2, F7, F3, Fz, F4, F8, T3, C3, Cz, C4, T4, T5, P3, Pz, P4, T6, O1, Oz, O2. The ground electrode was located in the AFz position, the reference ones were on the earlobes (positions A1, A2). A monopolar montage regarding the ear electrodes A1 and A2 was used. The impedance was maintained at a level below 5 kOhm. The video was recorded using a standard web camera.

**Part 2. Offline classifier training and 10-fold cross-validation test of classification accuracy**

After the first session with each subject, we got an EEG recording (with synchronized video) to be used as a calibration file in the offline classifier training. It was then marked as described earlier. Further, by passing this file through a sequence of OpenVibe scenarios, we trained a prediction model to be used in the next sessions online. Generally, we had to determine three components for classification: a) the selected frequency band for band-pass filtering of the EEG signal, and depending on it, both b) coefficients of spatial filters, and c) the configuration file of the trained model containing relevant coefficients for the online classification process. We had to determine what frequency band could give us the models with the most predictive power. The wide band (1.5-45 Hz) has been tested, and three narrower bands were also tested: 1.5-8 Hz, 1.5-14 Hz, and 4-15 Hz.

To increase the classification accuracy, we used spatial filtering of signals with the Common Spatial Pattern (CSP) method (Aydemir, 2016; Bird et al., 2019; Blankertz et al., 2008; Ramoser et al., 2001). Scenario-I (see Figure I below) allowed us to calculate the spatial filter coefficients. To accomplish this, epochs of 4 s were allocated around each event label so that the first 0.5 s of the epoch preceded the label, and the remaining 3.5 s followed it. The Temporal filter box used the band-pass filter by the Butterworth method, the filter order was 5. The selected dimension in the CSP Spatial Filter Trainer box was 12.

Figure I. Scenario-I in OpenVibe Designer calculates the spatial filter coefficients by the CSP algorithm. Each box contains its short functionality description on the left
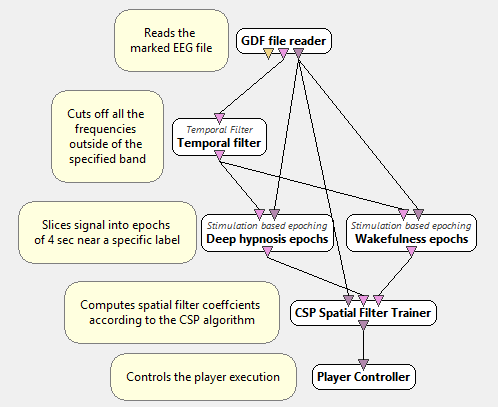

The obtained configuration files containing the spatial filter coefficients (component "b") were used in the following scenario (Scenario-II, see Figure II below), which is designed for classifier training. In Scenario-II, after frequency and spatial filtering, the signal was divided into two groups of epochs as in the previous scenario. Further, the feature extraction was performed for each of these two groups: the signal was split into blocks of 4 s every 0.5 s, and then the logarithmic band power was computed as log (1+x) where "x" is the mean of the squares of the signal in each of the blocks. This value was subsequently converted into a feature vector, which then went to the Classifier Trainer box. Thus, the Classifier Trainer was receiving two types of feature vectors (corresponding to the state "W" and the state "D"), and after the end of the training, it produced a configuration file containing the relevant coefficients of the trained model (component "c") to be used during online sessions in the next scenario. Linear Discriminant Analysis (LDA) was applied as an algorithm to classify two types of feature vectors.

The usage of the proposed method in the therapeutic context assumes that EEG recordings of the first session will serve as a calibration file, but for research purposes, we marked up all EEGs and ran them through Scenarios-I and -II to calculate the classification accuracy for each session. The Classifier Trainer box executed a 10-fold cross-validation test for four types of our models to estimate what frequency band provided the most accurate average results. This method assumes testing the predictive power of the model based on the same data that were used for the training.

Figure II. Scenario-II in OpenVibe Designer. Each box contains its short functionality description on the left, except those already described in Scenario-I
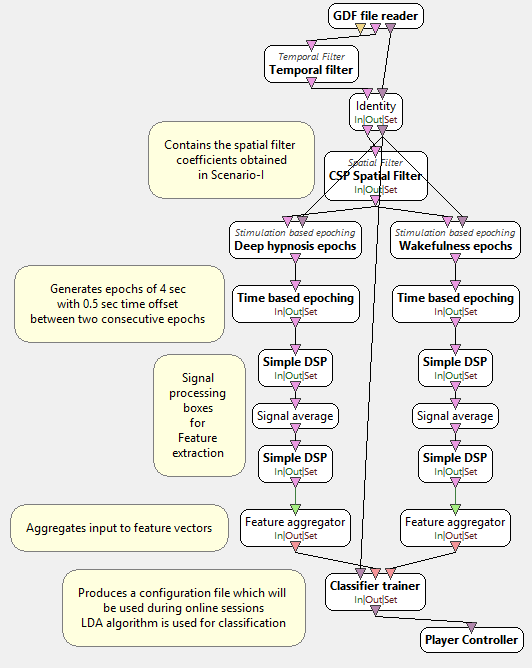

**Part 3. Testing the trained models in real time**

All three obtained components ("a", "b" and "c") were applied in the second and the following sessions to provide the online classification process. In Scenario-III (see Figure III below), the real-time EEG signal underwent band-pass and spatial filtering and transformed to the feature vectors like in Scenario-II. The feature vectors were then sent to the Classifier Processor box, containing the configuration file of our trained model (component "c"). As a result, the system was able to predict which of the two states and with what probability the patient was in at a moment, which was providing Probability Values as an output parameter. This output is a matrix of interdependent probabilities of matching current EEG patterns to one state and another. For visualization convenience, we selected from the matrix only the probability of deep hypnosis and displayed this value fluctuating from 0 to 1 online using the Continuous Oscilloscope box. At that point, the closer to "1", the more probable the deepest stage and less probable the wakefulness. Supposedly, it could be interpreted as a depth level measuring, meaning the higher the probability, the deeper the hypnosis. Thus, Scenario-III was built to provide a single integrative individualized continuously changing parameter displayed in the form of a curve during the session. We called this curve the Predictive curve. For the convenience of displaying data over a large time interval, to smooth the curve we used the function Moving average (Immediate) in the Epoch average box.

At the baseline EEG registration of each second and subsequent session by alternating testing, we selected the frequency band and other corresponding components for the trained model that predicted the minimal deep hypnosis probability. This was the signal that chosen components at least were properly predicting the wakefulness state. Further, we conducted an online session with them to watch the Predictive curve.

Figure III. Scenario-III in OpenVibe Designer. Each box contains its short functionality description on the left except those already described in the previous scenarios

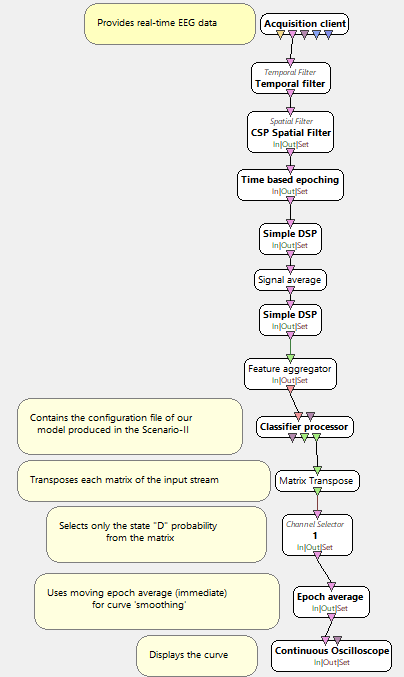

**Part 4. Estimating the accuracy of the models based on new data (data of the second and subsequent sessions)**

The obtained EEG recordings of the second and subsequent sessions were labeled according to the same principle used for the first (calibration) sessions. To eliminate bias, this was done by a specialist not privy to the predictive outcomes of these sessions. Next, the marked file was passed through Scenario-IV (see Figure IV below), which was almost completely the same as Scenario-III, but this labeled EEG was the data source in this case. Also, this scenario uses the Classifier Accuracy Measure box which computes the accuracy of the model given the results from the classifier, compared to the labels received. This is measured as the percentage of correctly classified epochs for the entire recording (Kohavi & Provost, 1998). At this stage of our analysis, we tested the accuracy of four types of models for each of the second and subsequent sessions.

The training on marked-up data of each second and subsequent session resulted in obtaining their own, auxiliary classification models. We did not use them for further online sessions, but we put them into Scenario-IV and passed the same data, on which these models were trained, through this scenario. This way, after each visit, we built a curve that reflected the dynamics of the hypnotic depth in this session as it really was, i.e., with maximal accuracy. We called it the Native curve. Then, the configurations of the Native and Predictive curves of the same sessions were visually compared to additionally assess how accurately the model trained on the first session data was able to reflect the actual picture of the subsequent ones.

Figure IV. Scenario-IV in OpenVibe Designer. Each box contains its short functionality description on the left except those already described in the previous scenarios
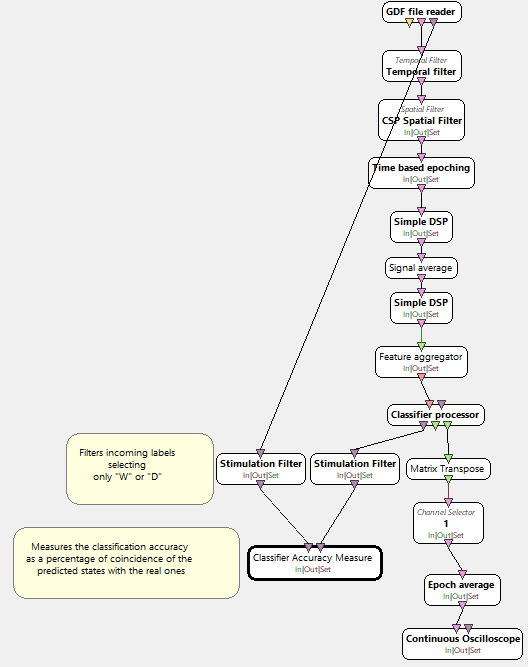
