## Supplementary materials B for "Teaching a computer to assess hypnotic depth: A pilot study"

Topographic maps displaying the changes in the power of different rhythms for different localisations after achieving deep hypnosis. Changes are displayed in color according to the graduation of a nearby color scale. The color scale is in μV². The data for all included patients and sessions are presented

**Patient A, session 1**
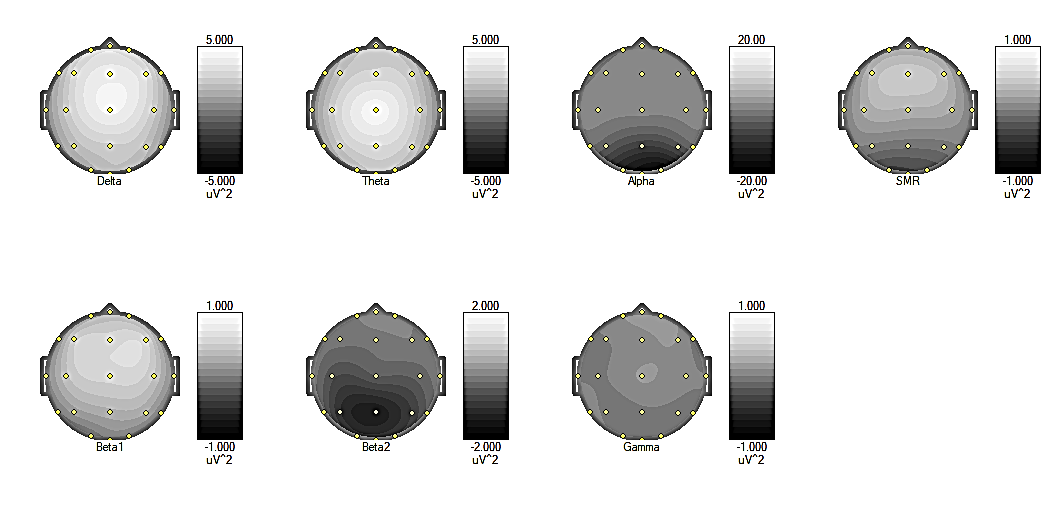

**Patient A, session 2**

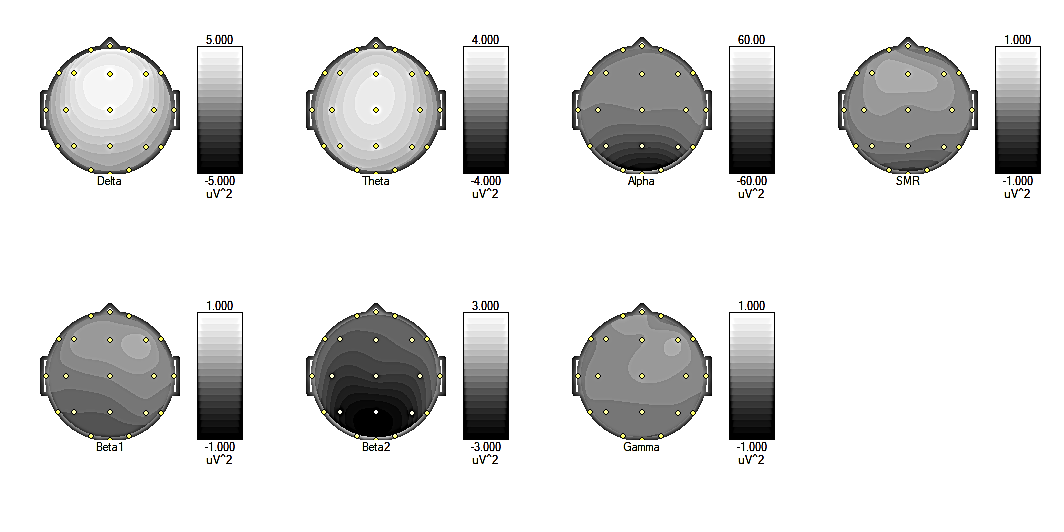

**Patient A, session 3**

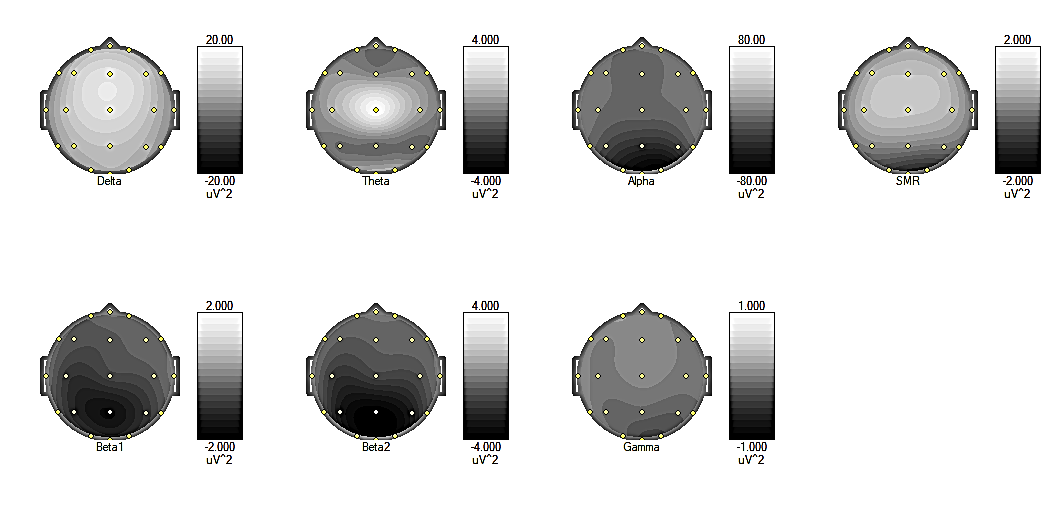

**Patient A, session 4**

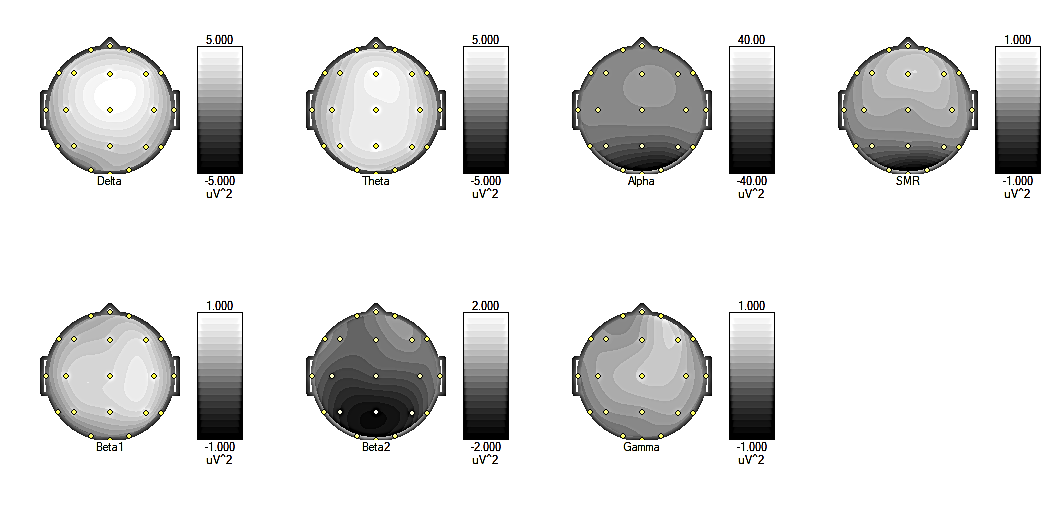

**Patient A, session 5**

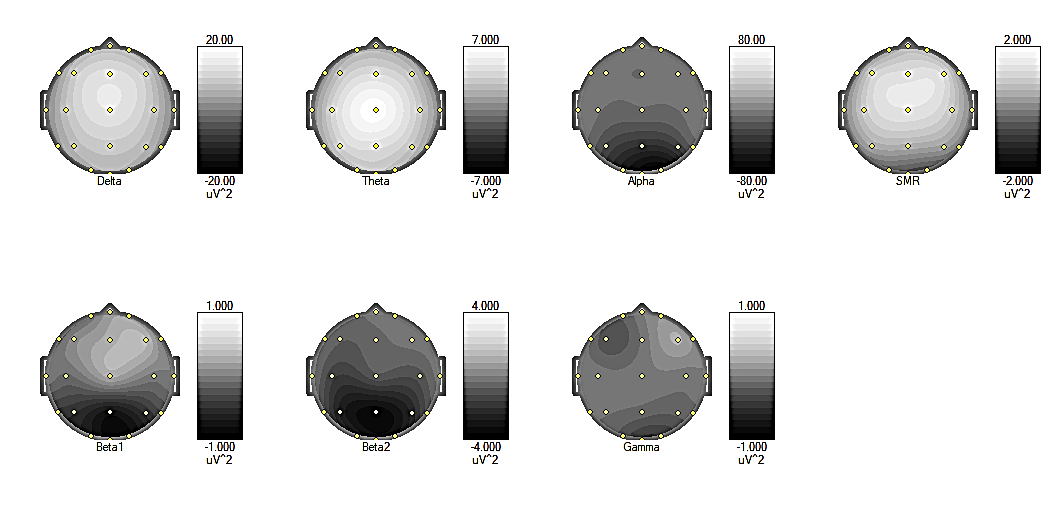

**Patient A, session 6**

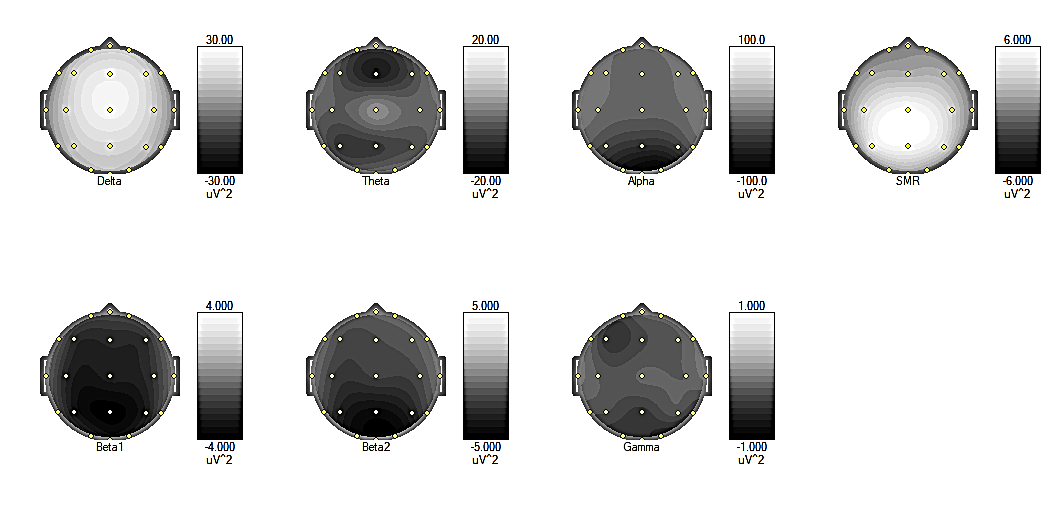

**Patient A, session 7**

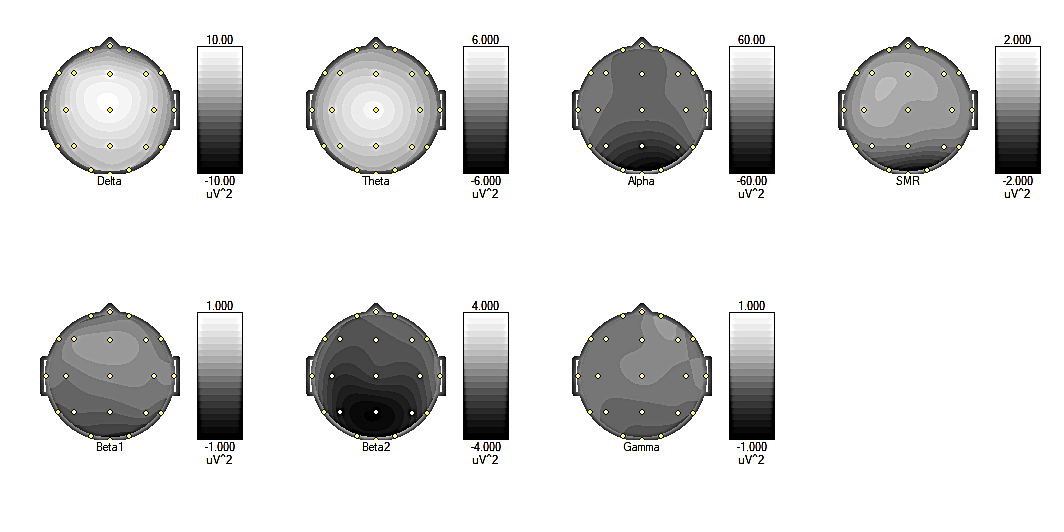

**Patient E, session 1**

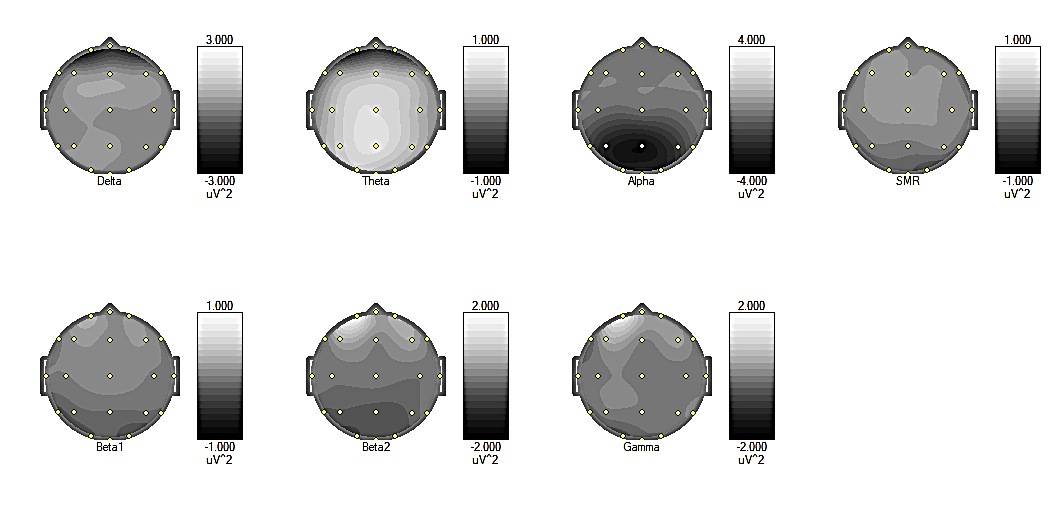

**Patient E, session 2**

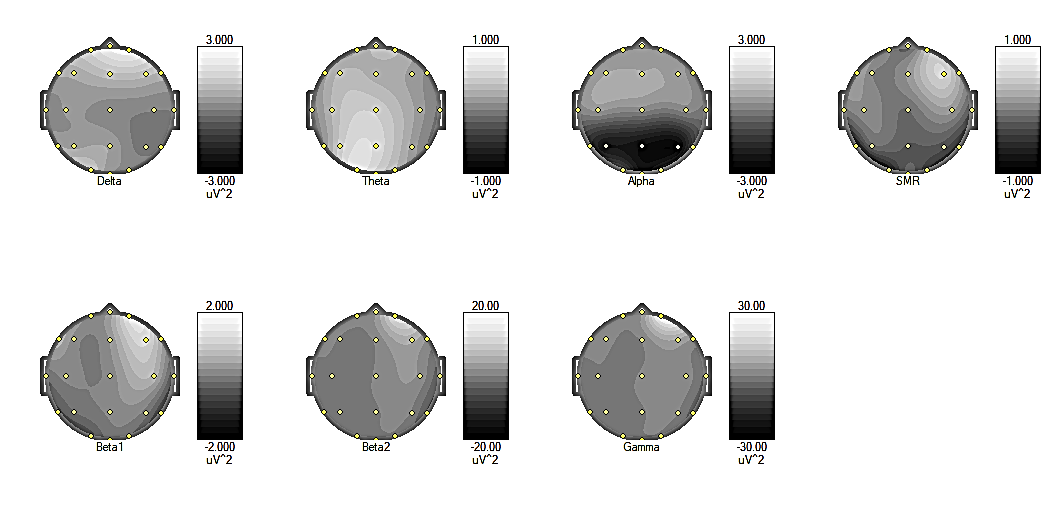

**Patient E, session 3**

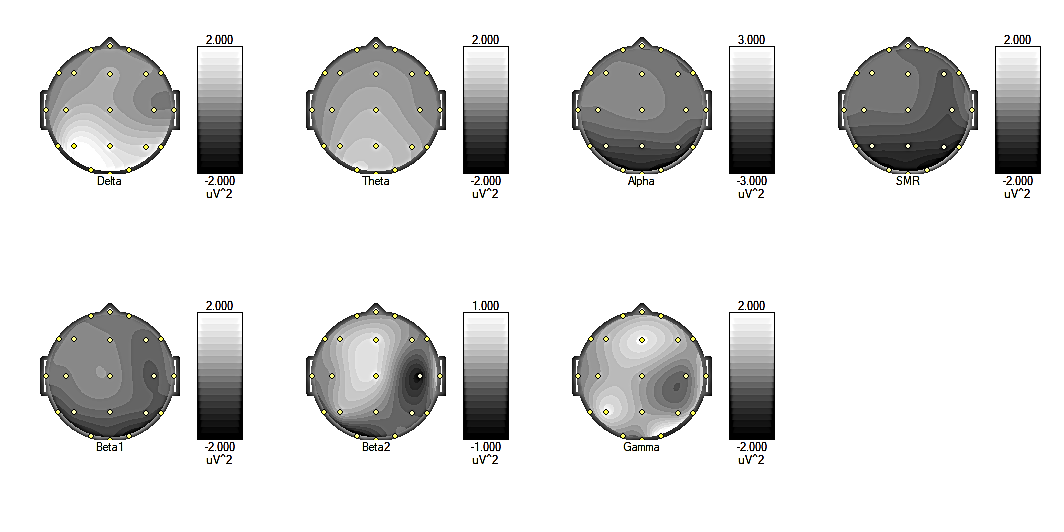

**Patient E, session 4**

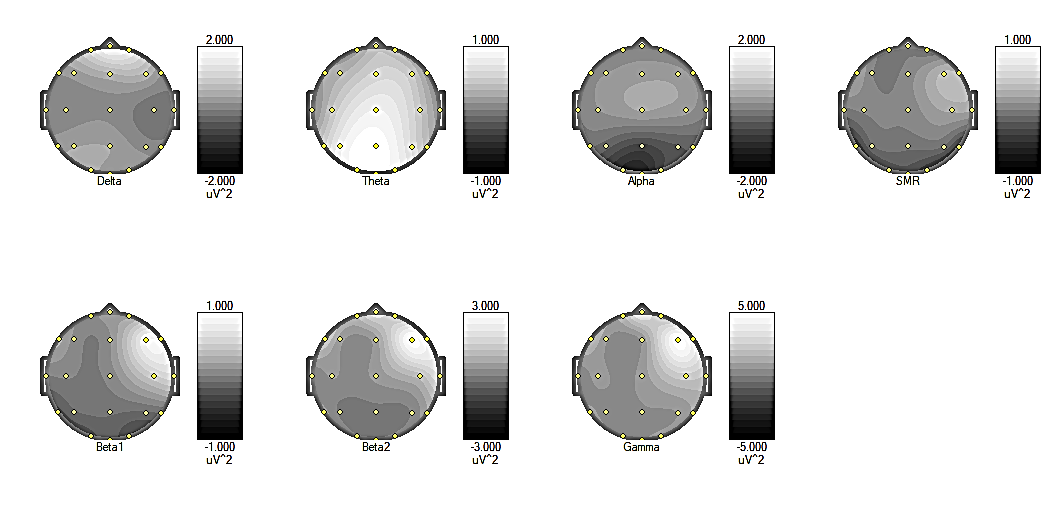

**Patient E, session 6 (the 5^th^ was excluded)**

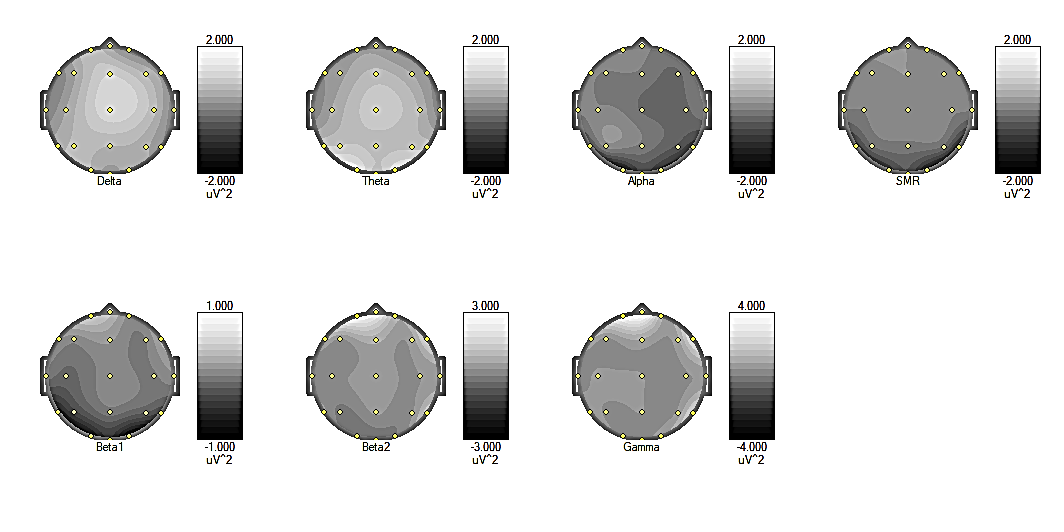

**Patient G, session 1**
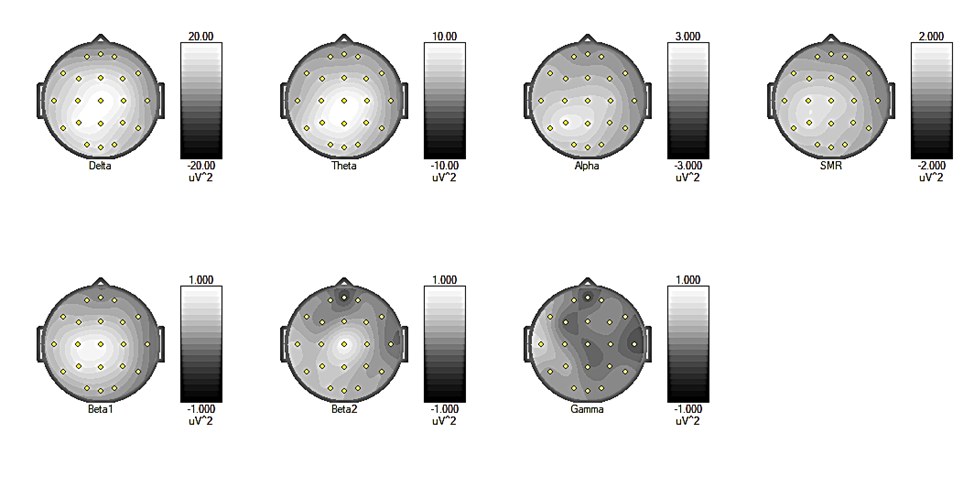

**Patient G, session 2**
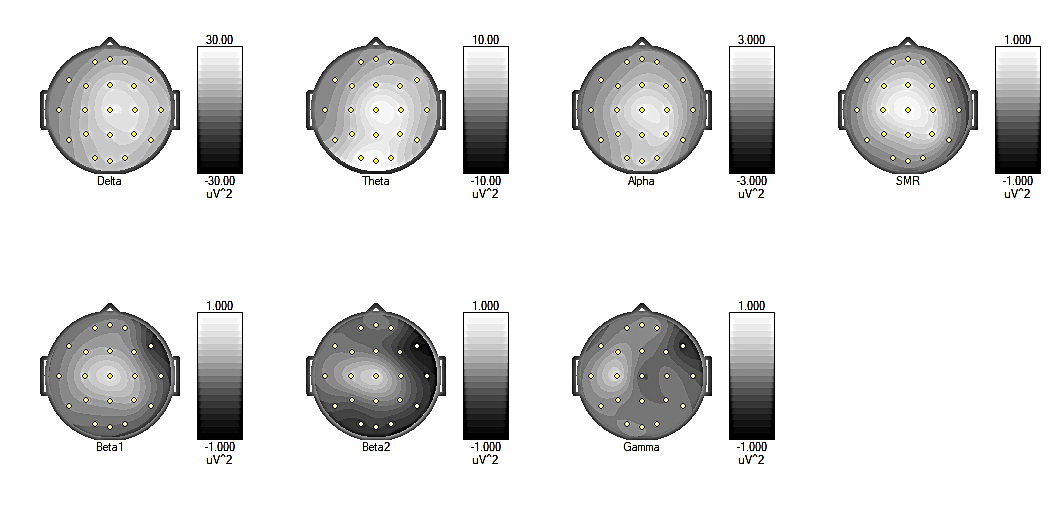

**Patient G, session 3**
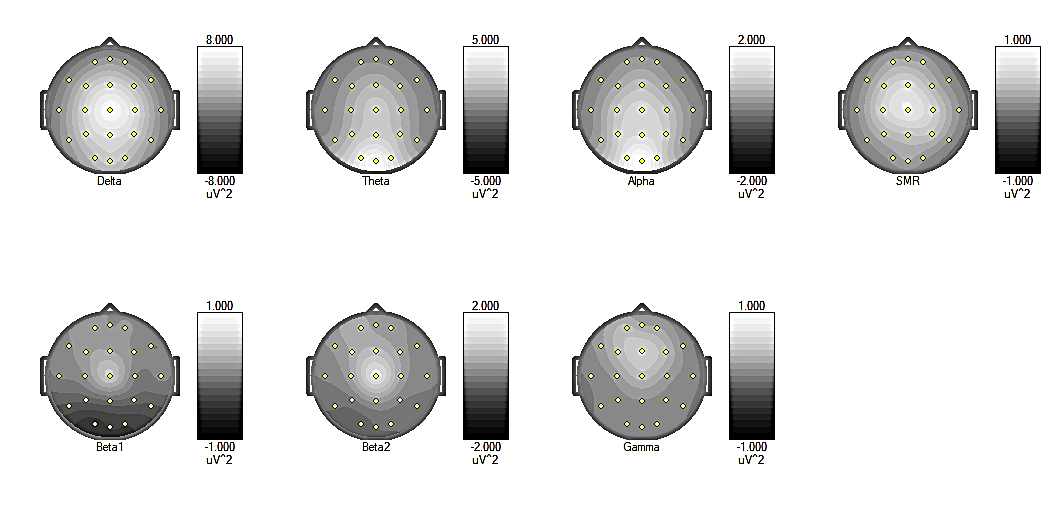

**Patient G, session 4**
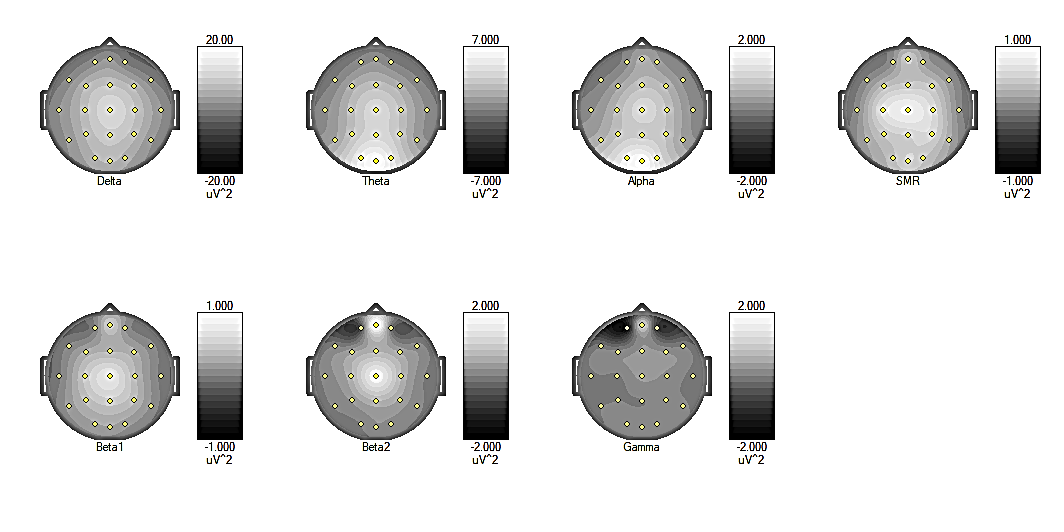

**Patient S, session 1**
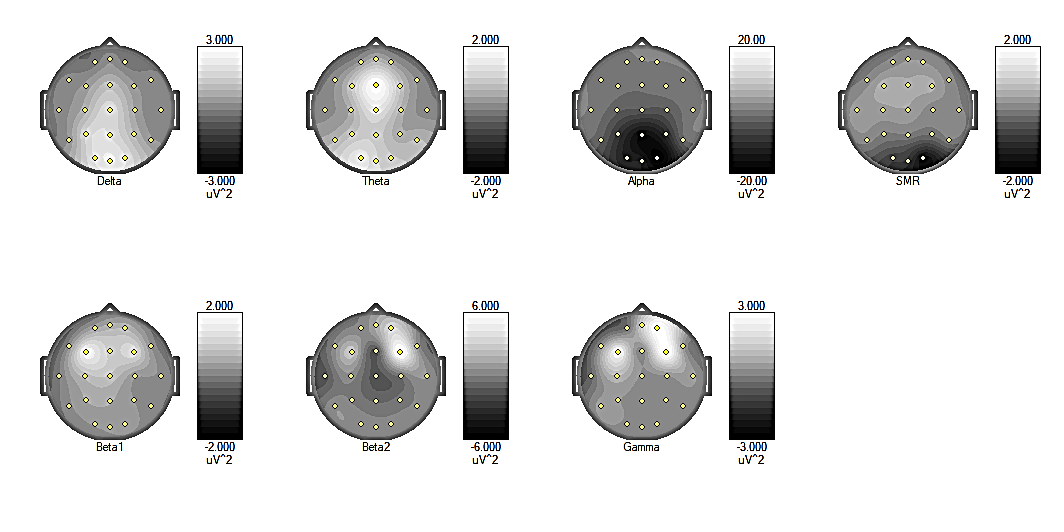

**Patient S, session 2**

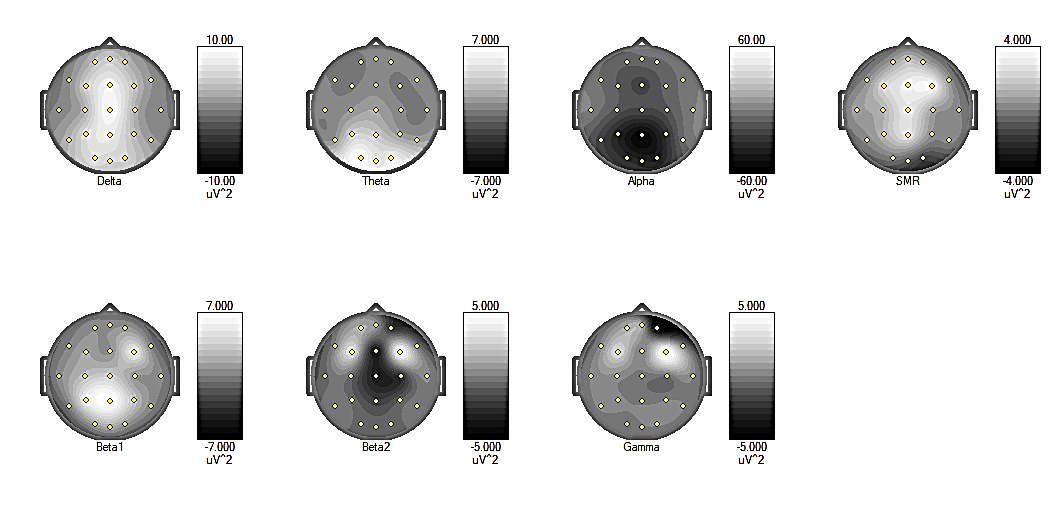

**Patient S, session 3**
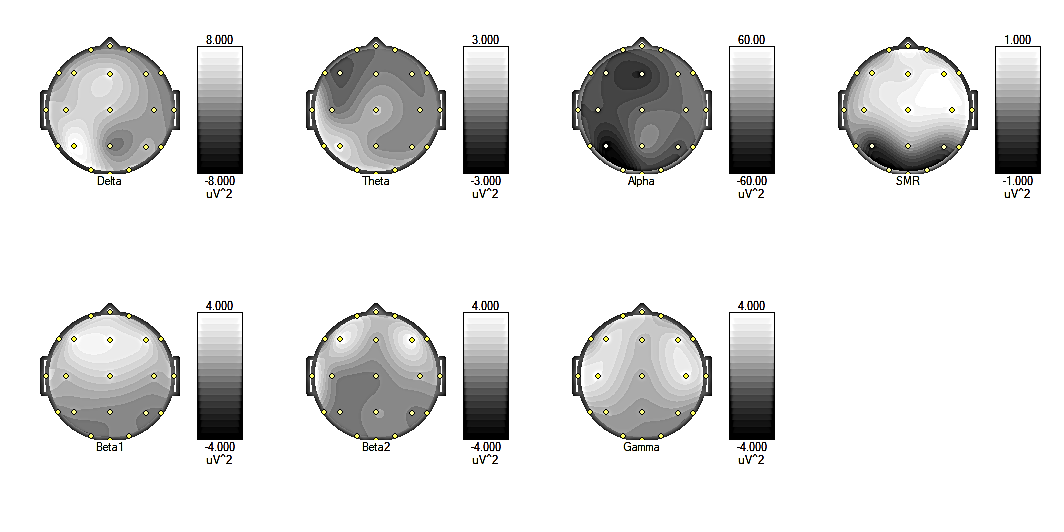

**Patient O, session 1**
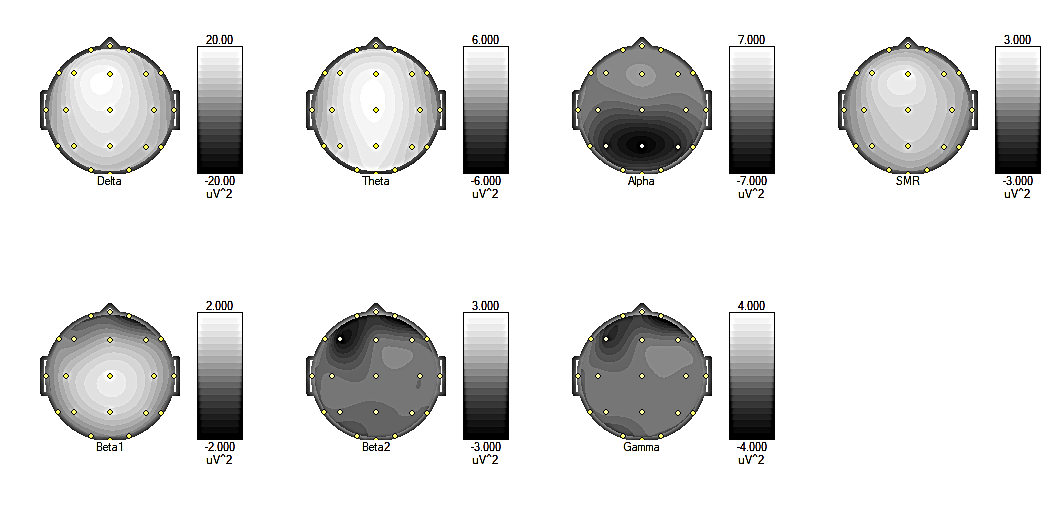

**Patient O, session 2**
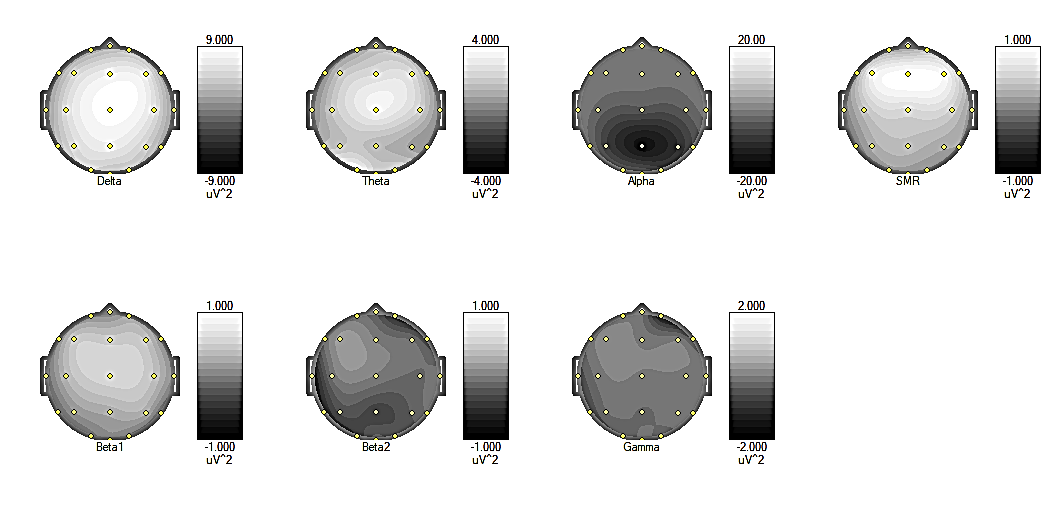

**Patient N, session 1**
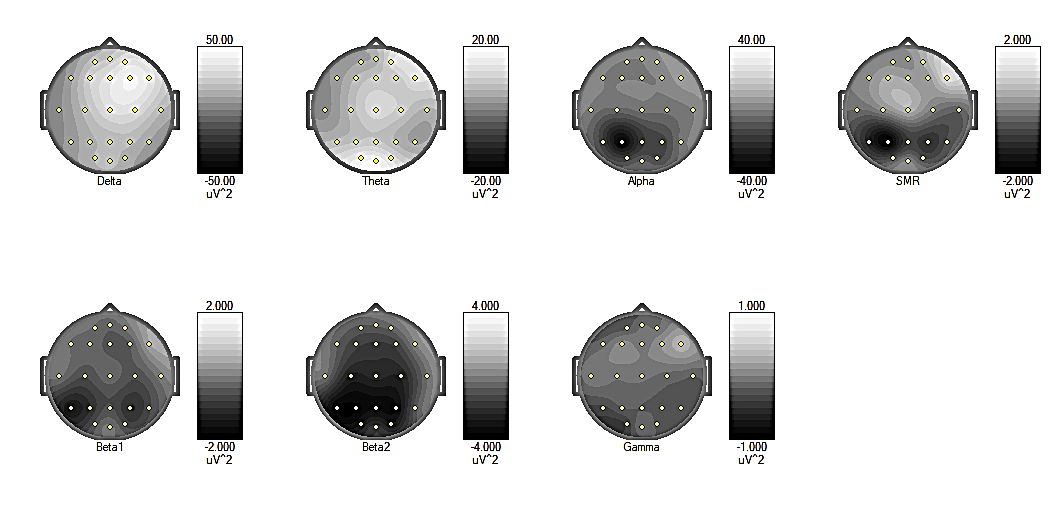

**Patient N, session 2**
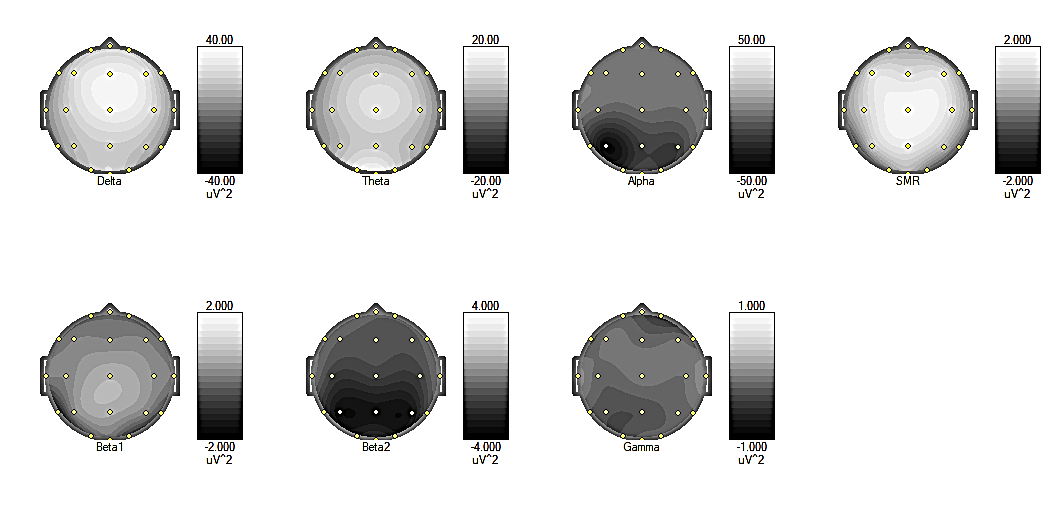

**Patient V, session 1**

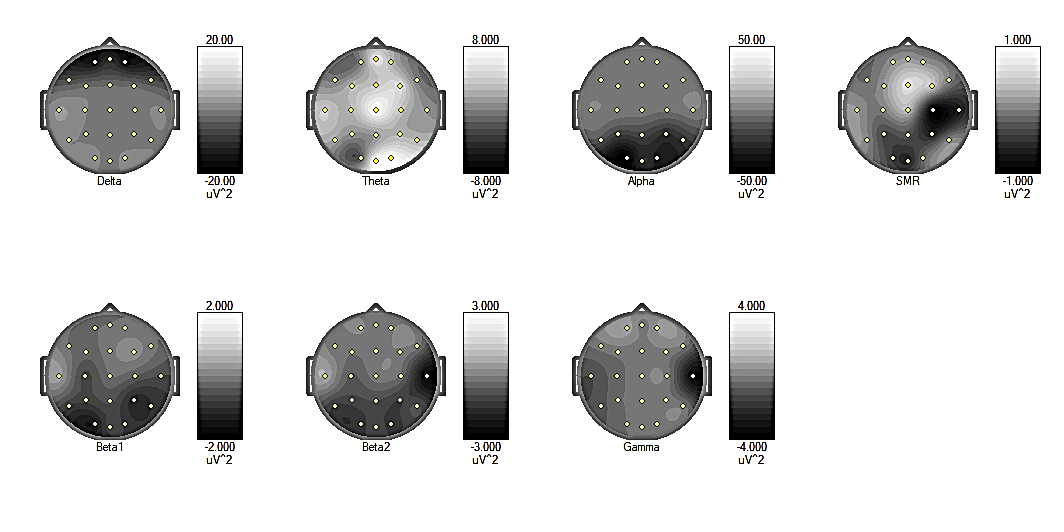

**Patient V, session 2**
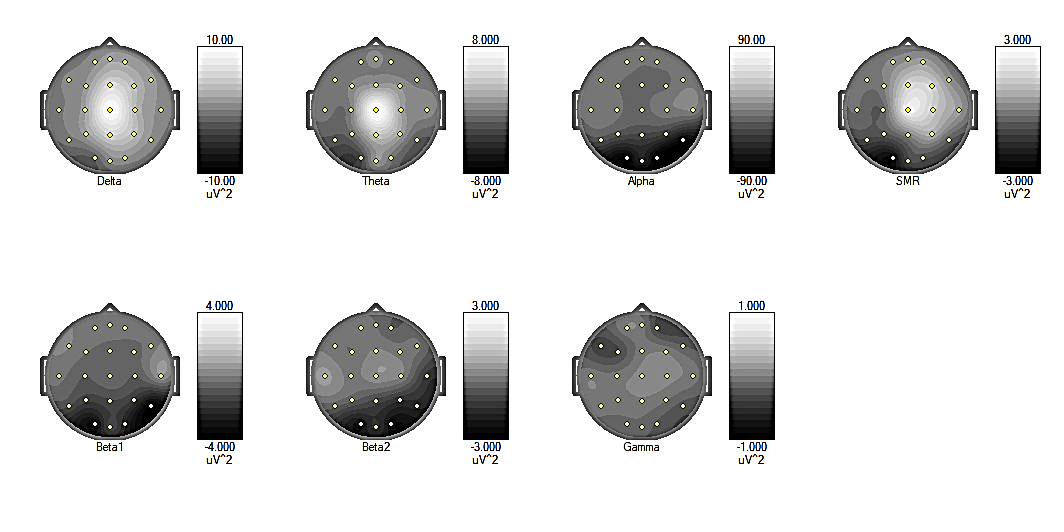

**Patient C, session 1**
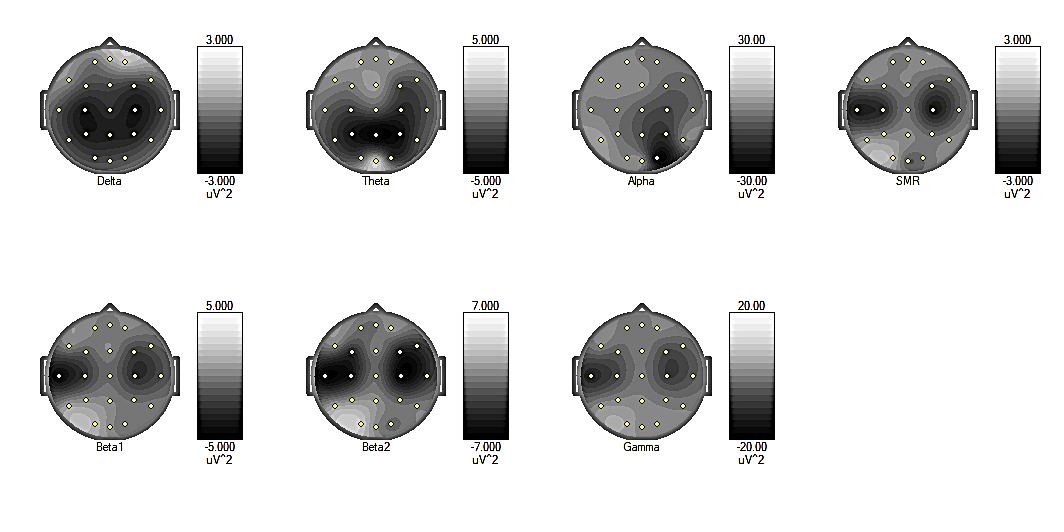

**Patient C, session 2**
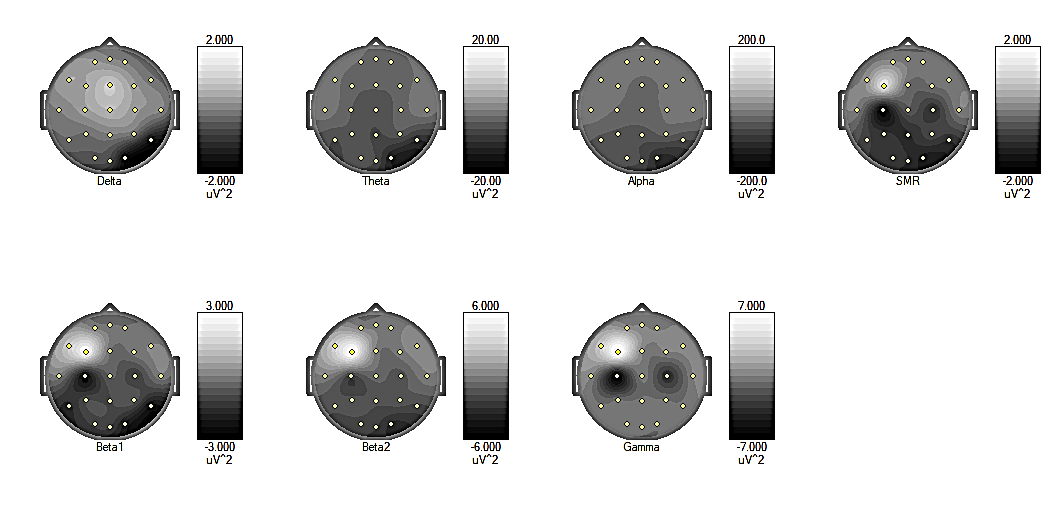
