## Supplementary materials C for "Teaching a computer to assess hypnotic depth: A pilot study"

**The pairs of the Predictive (upper) and Native (lower) curves for each patient in the study for the second and the subsequent sessions**

The frequency band used to build the Predictive and Native curves is indicated for each patient. The band that provided a high average classification accuracy for each exact patient was selected.

For the convenience of displaying data over a large time interval, the curves are smoothed using the Moving epoch average (Immediate) function in the Epoch average box. The number of 4-s epochs with an overlap of 0.5 s used for averaging is indicated for each pair.

**Patient A**, the band used is 1.5-14 Hz

The 2^nd^ session: the number of epochs for averaging is 100

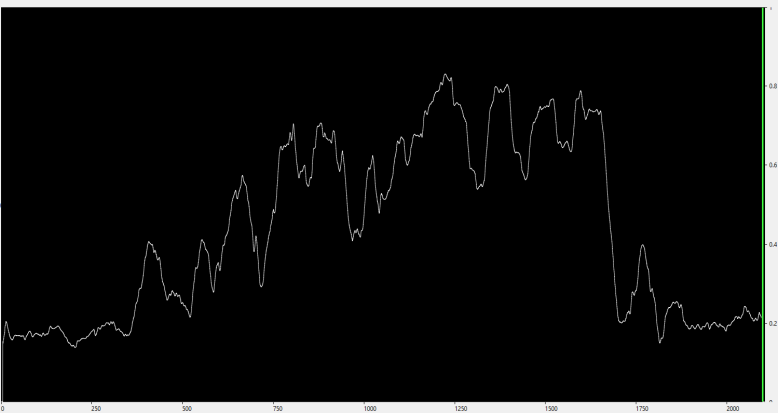

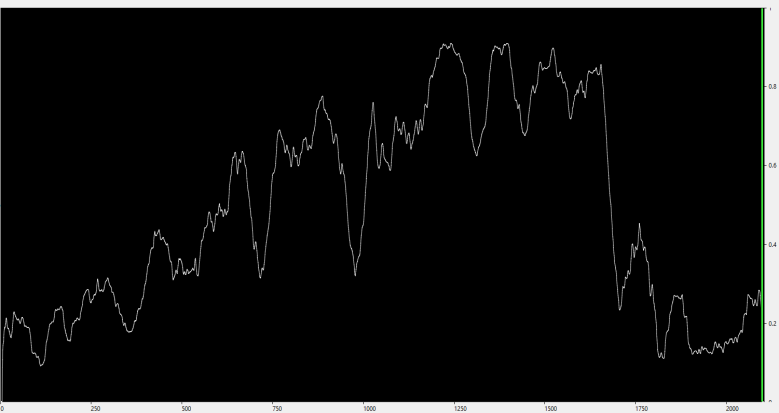

The 3^rd^ session: the number of epochs for averaging is 50

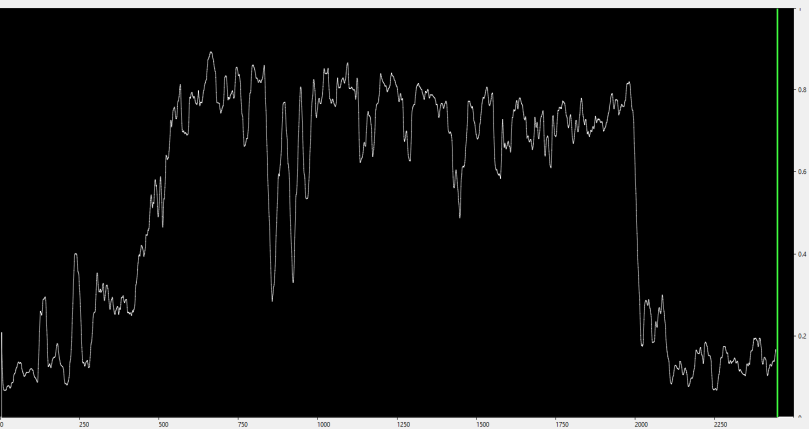

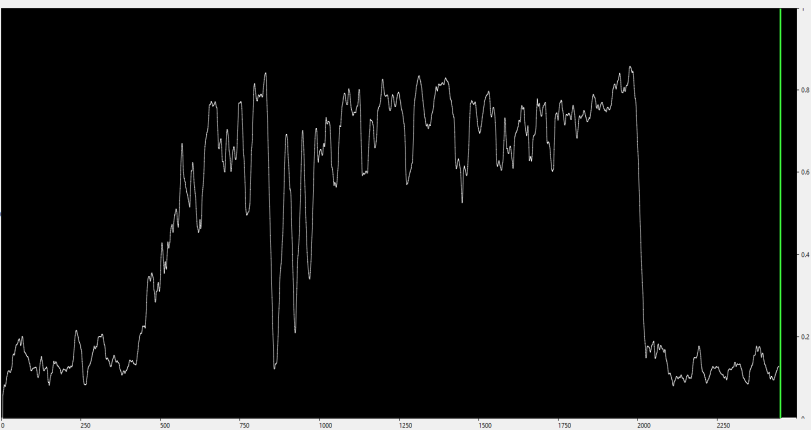

The 4^th^ session: the number of epochs for averaging is 100

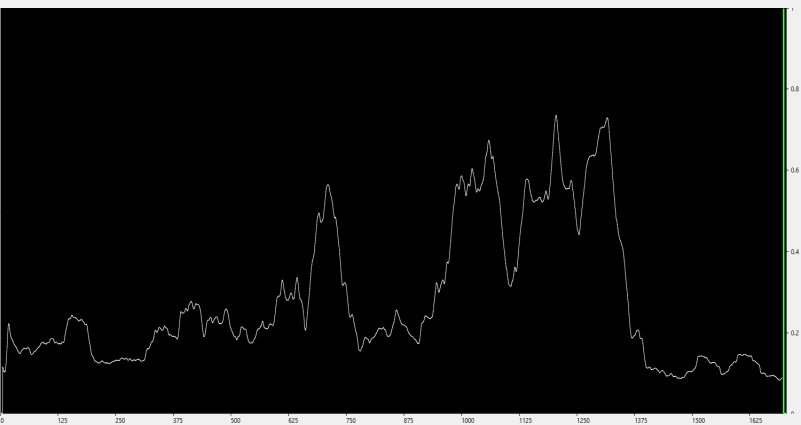

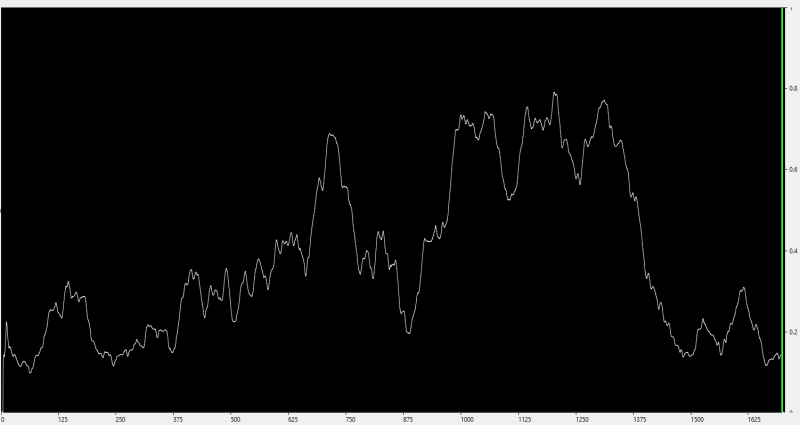

The 5^th^ session: the number of epochs for averaging is 50

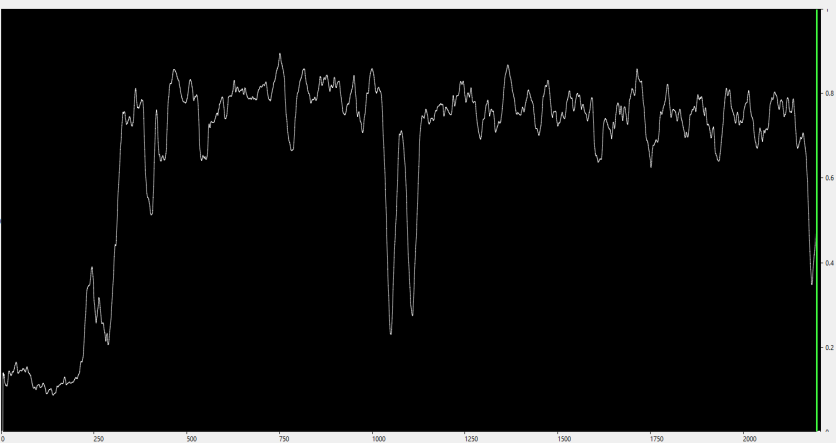

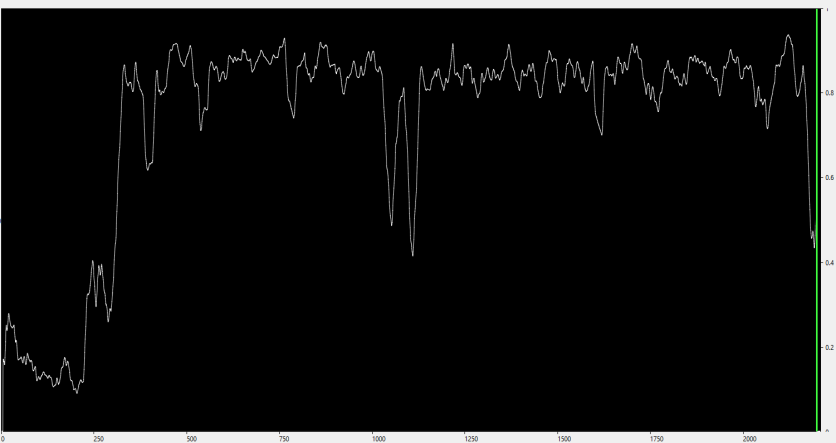

The 6^th^ session: the number of epochs for averaging is 50

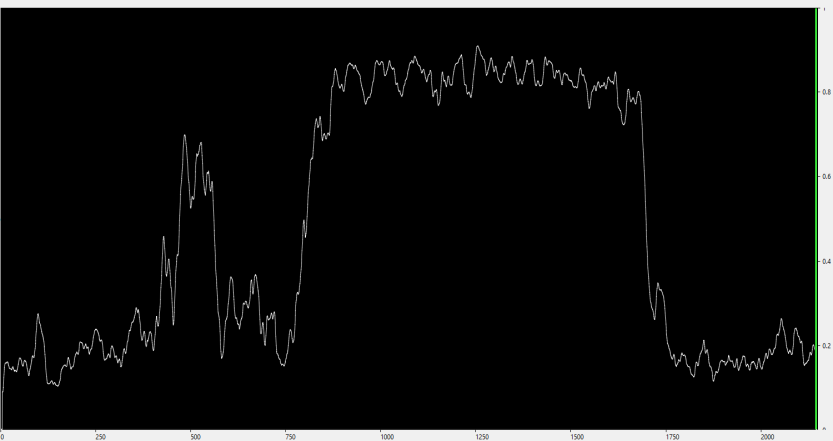

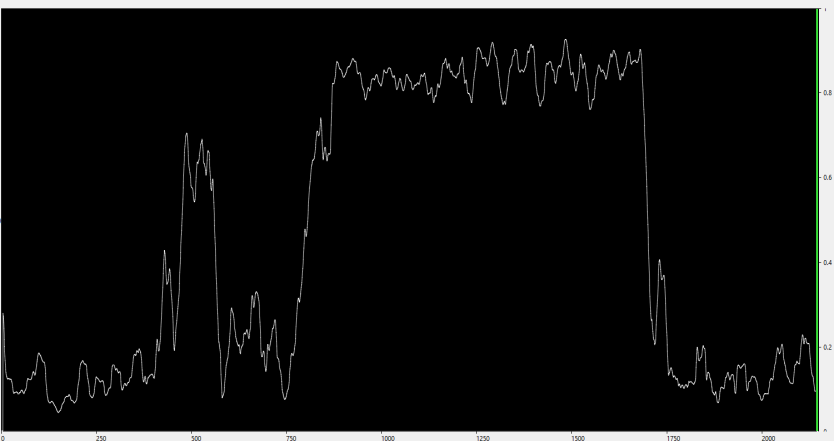

The pair of curves for the 7^th^ session of this patient is displayed in Figures 2 and 4 in the main text

**Patient E**, the band used is 4-15 Hz

The 2^nd^ session: the number of epochs for averaging is 150

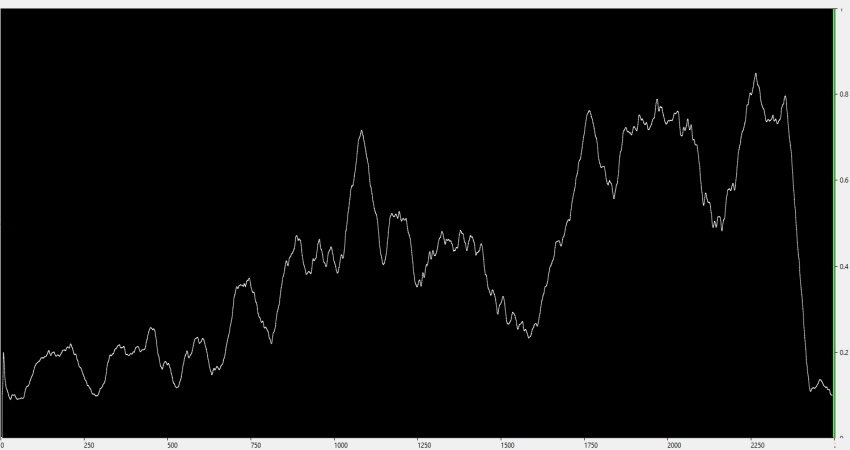

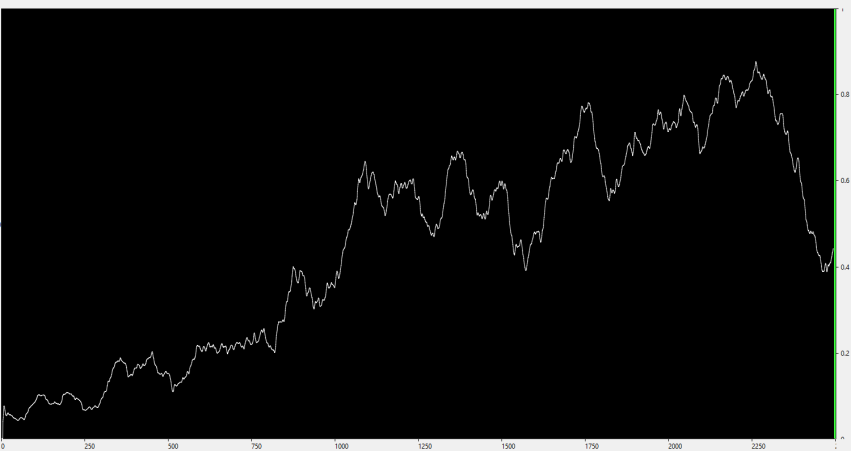

The 3^rd^ session: the number of epochs for averaging is 50

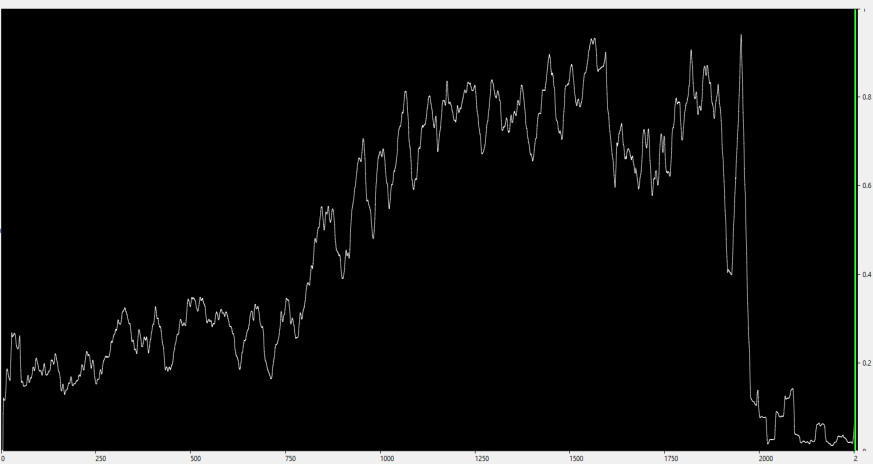

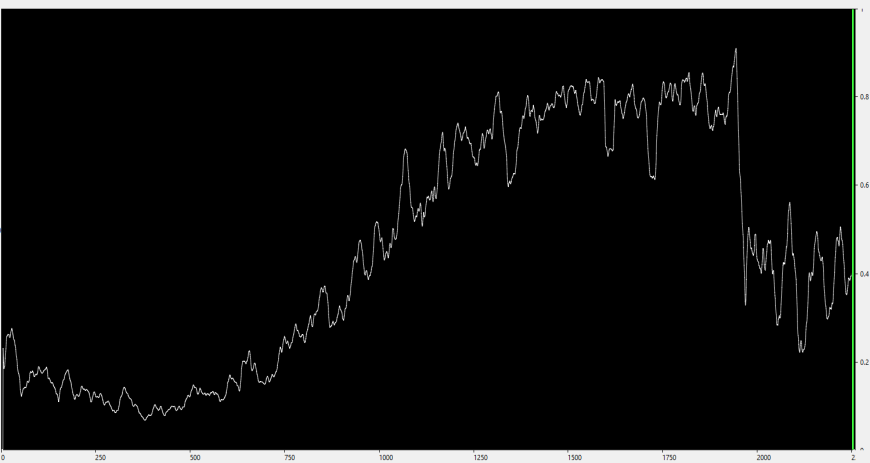

The 4^th^ session: the number of epochs for averaging is 150

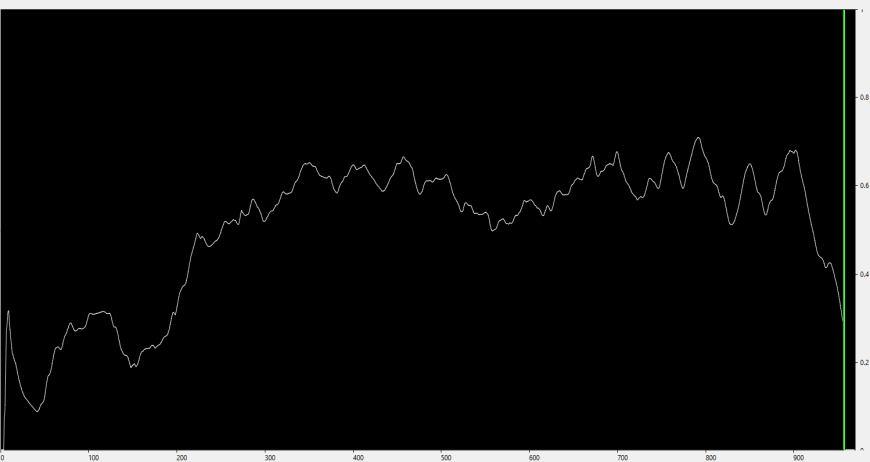

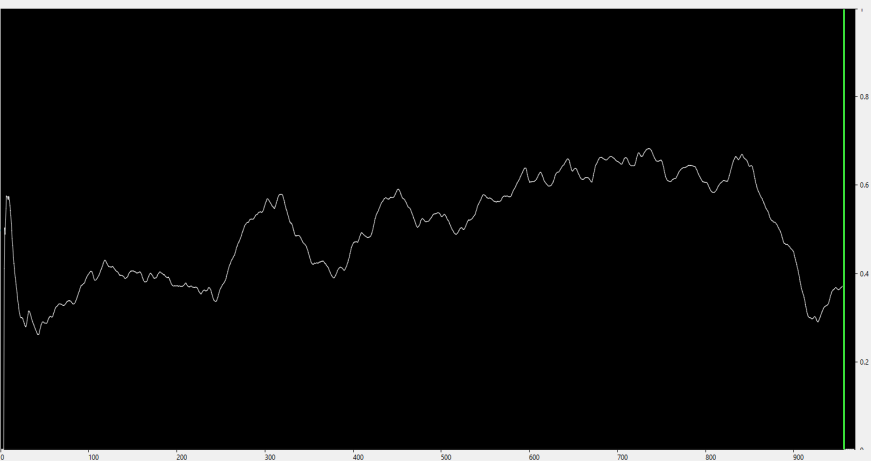

The 5^th^ session was excluded from the analysis

The 6^th^ session: the number of epochs for averaging is 100

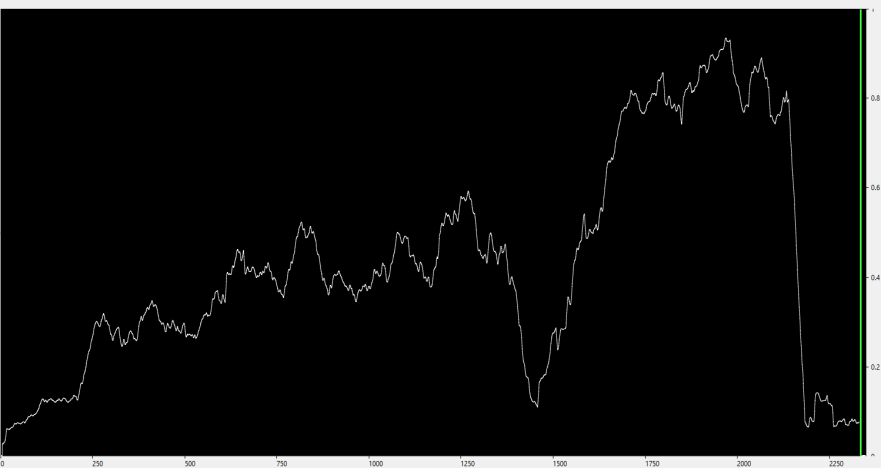

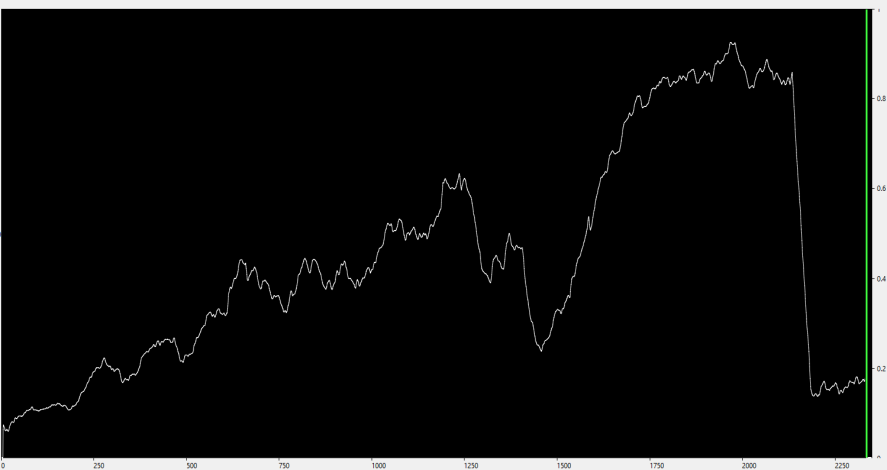

**Patient G**, the band used is 1.5-8 Hz

The 2^nd^ session: the number of epochs for averaging is 100

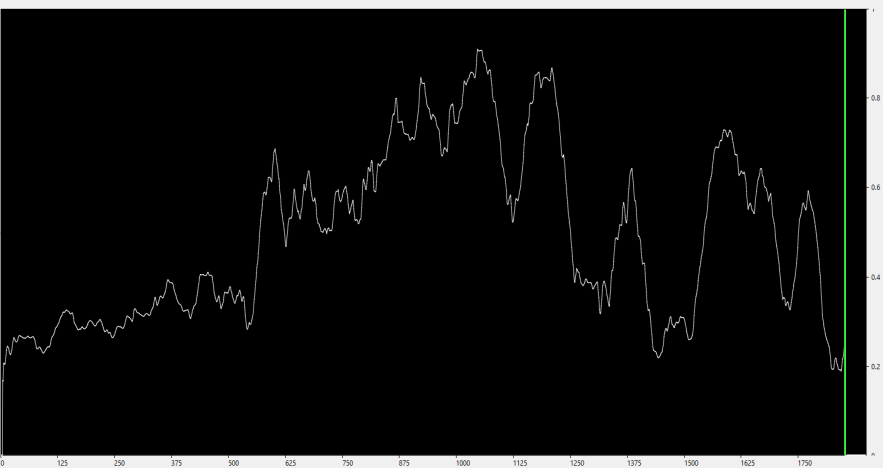

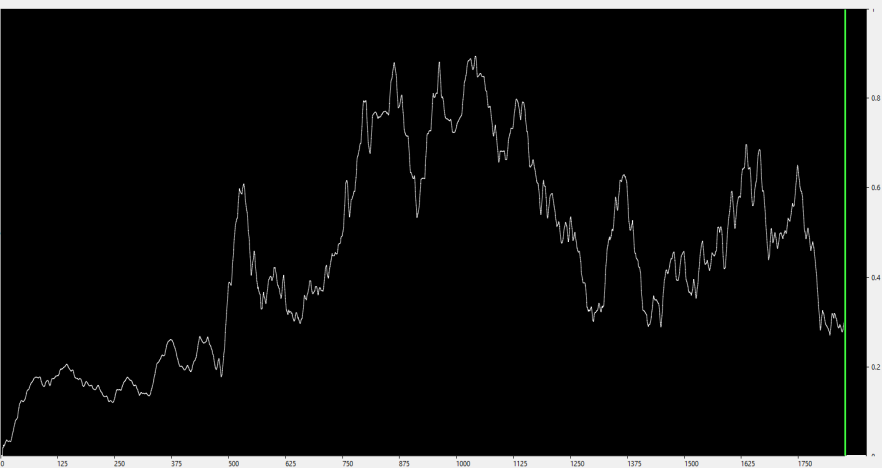

The 3^rd^ session: the number of epochs for averaging is 100

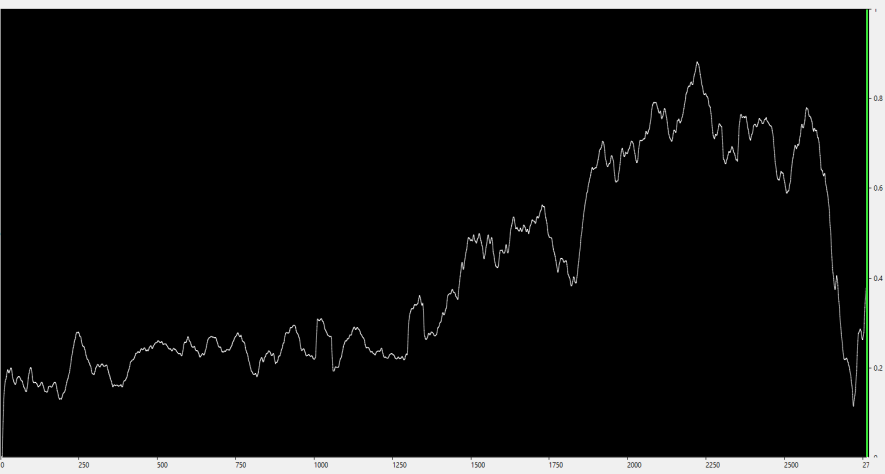

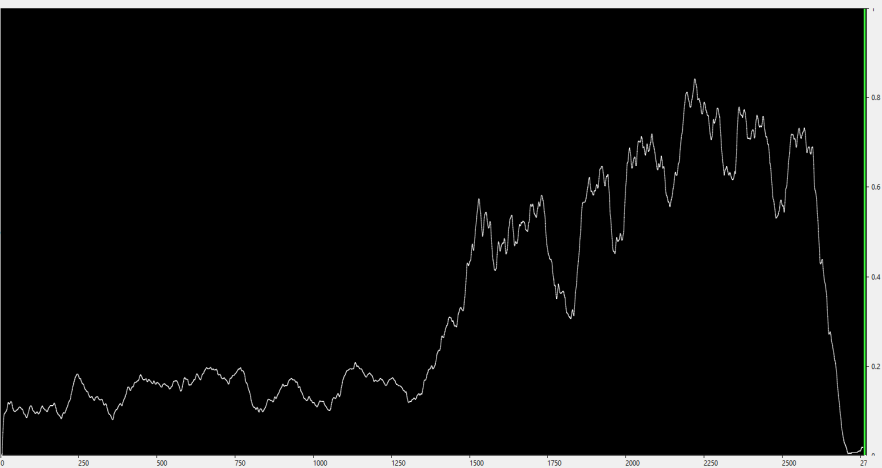

The 4^th^ session: the number of epochs for averaging is 100

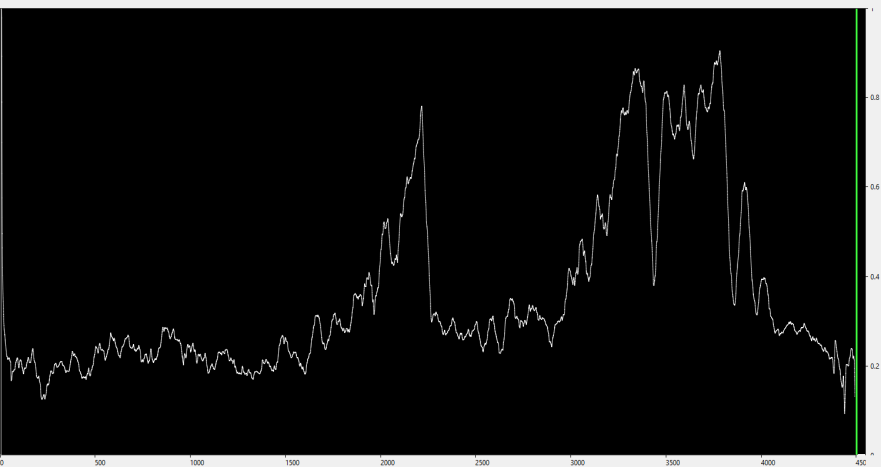

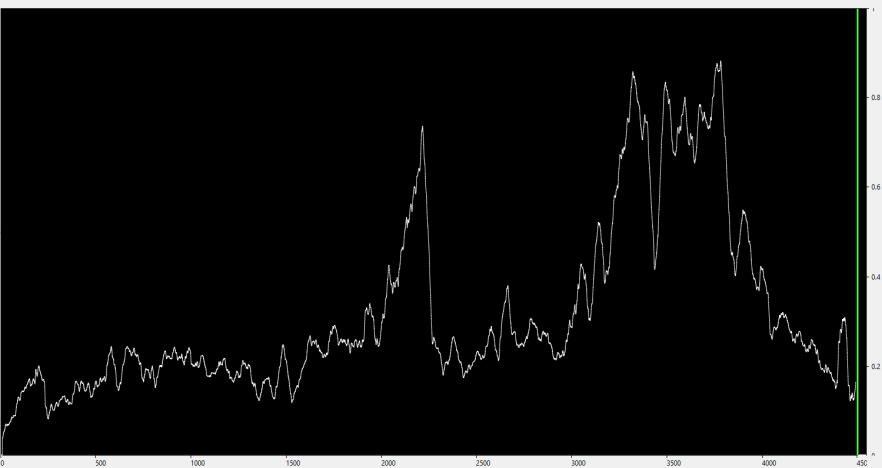

**Patient S**, the band used is 1.5-8 Hz

The 2^nd^ session: the number of epochs for averaging is 100

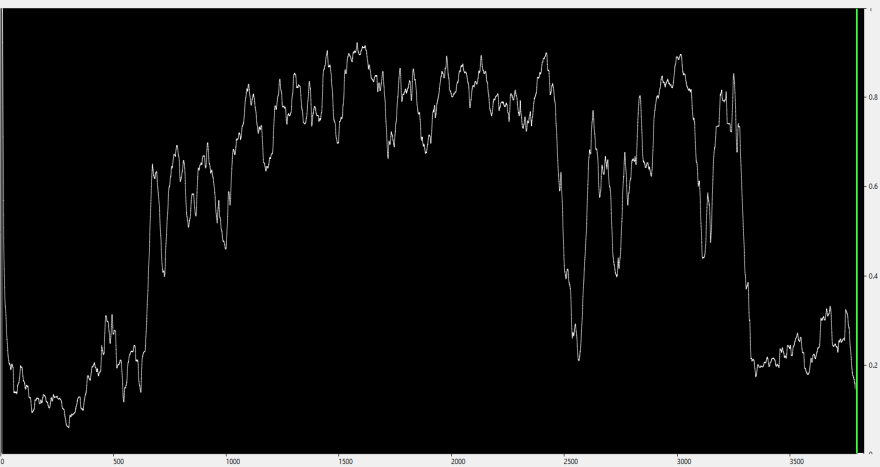

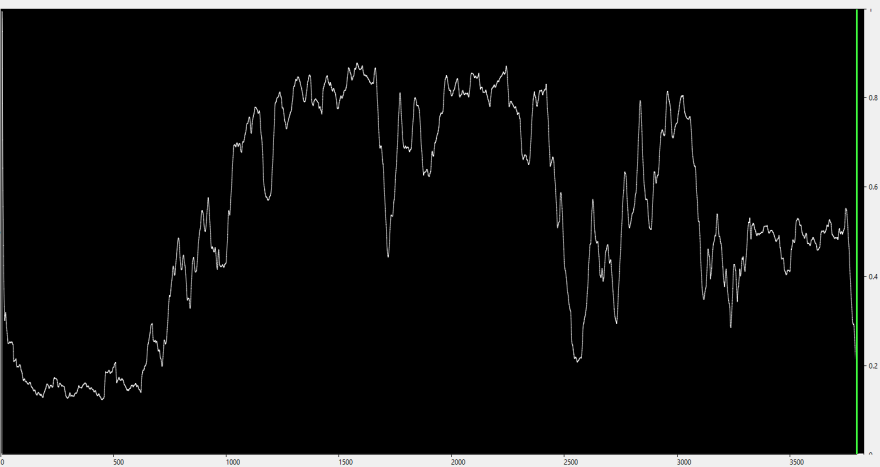

The 3^rd^ session: the number of epochs for averaging is 100

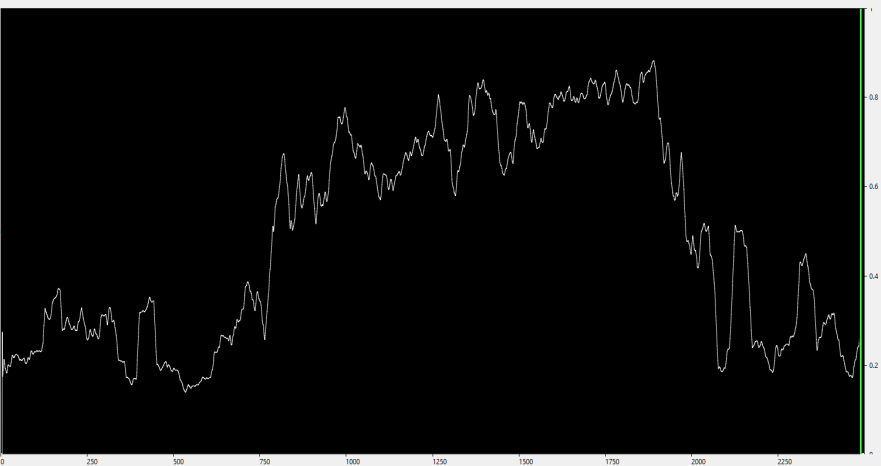

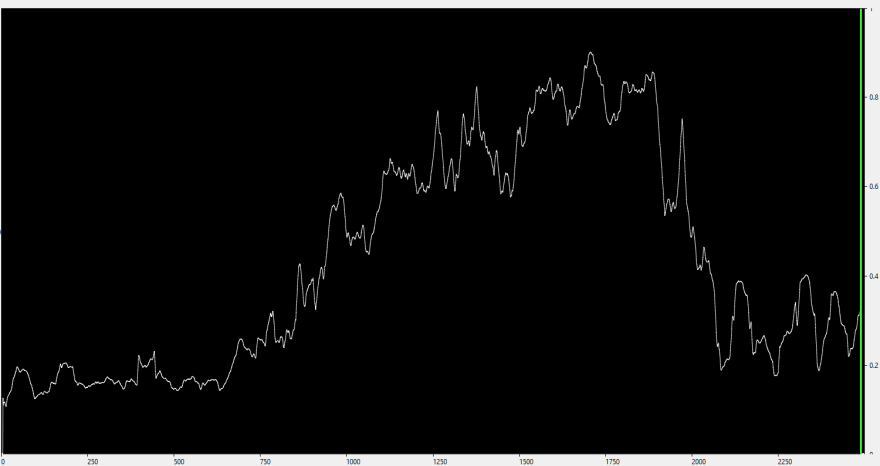

**Patient O**, the band used is 4-15 Hz

The 2^nd^ session: the number of epochs for averaging is 50

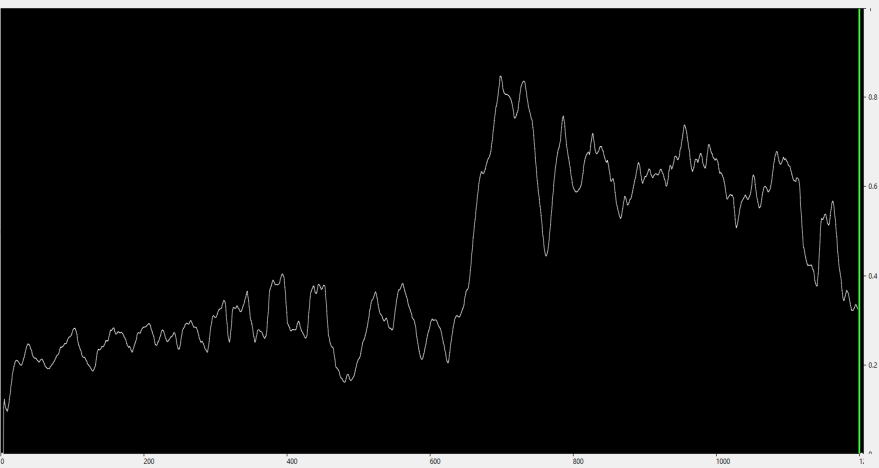

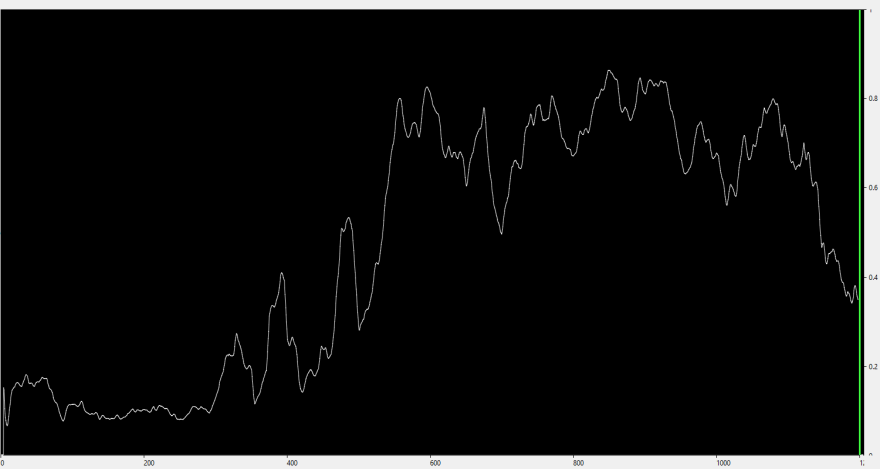

**Patient N**, the band used is 1.5-45 Hz

The 2^nd^ session: the number of epochs for averaging is 150

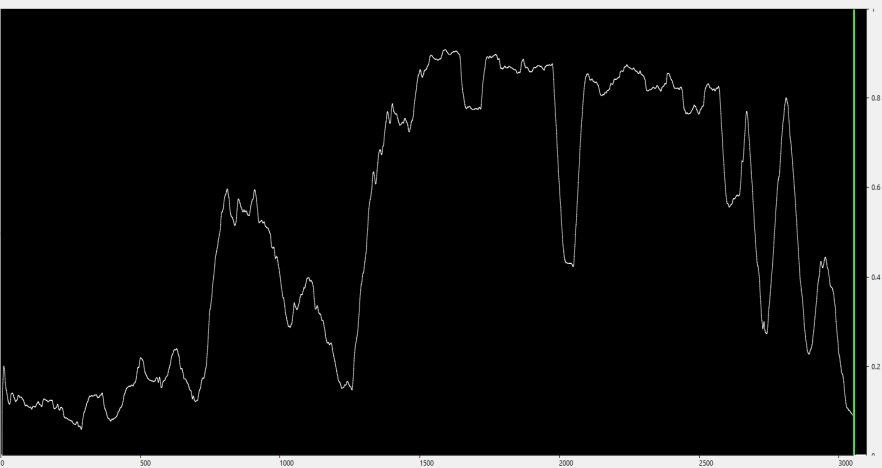

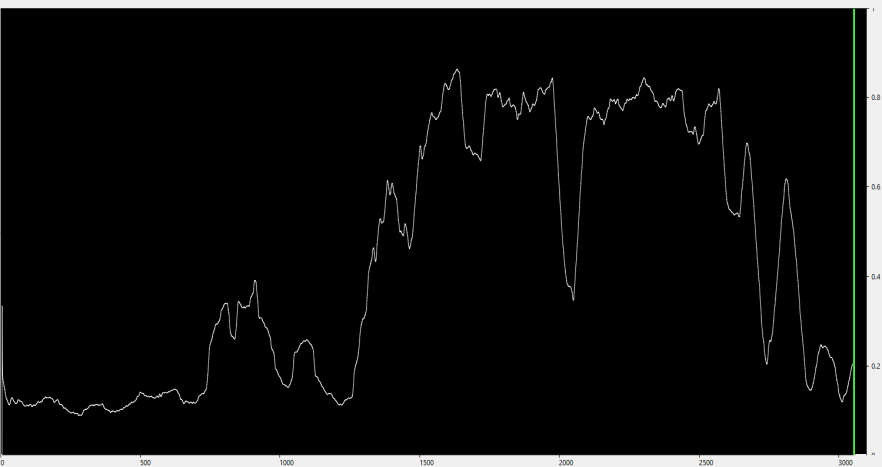

**Patient V**, the band used is 1.5-45 Hz

The 2^nd^ session: the number of epochs for averaging is 100

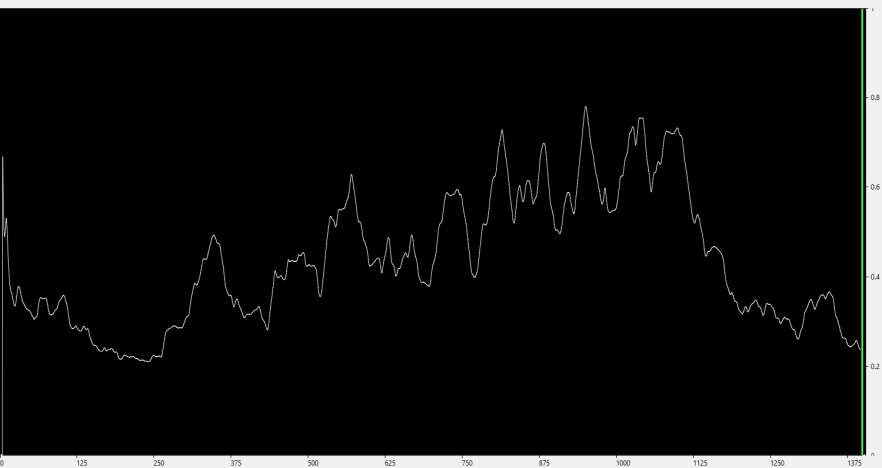

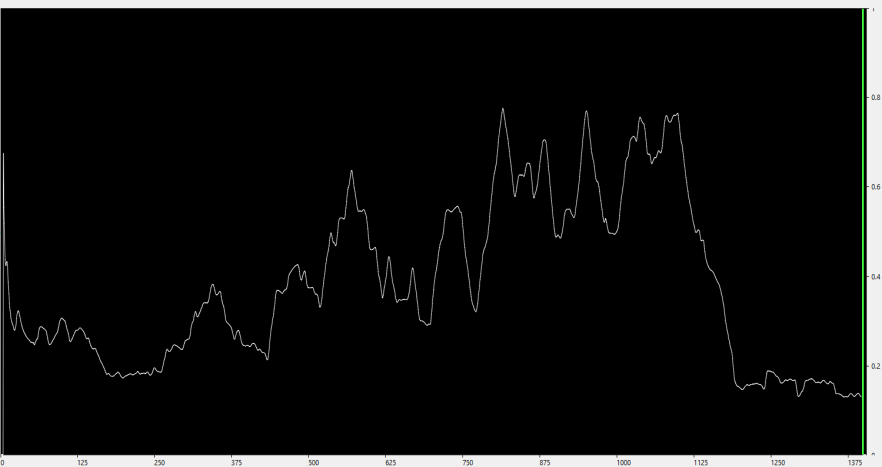

Patient C, the band used is 4-15 Hz

The 2^nd^ session: the number of epochs for averaging is 50

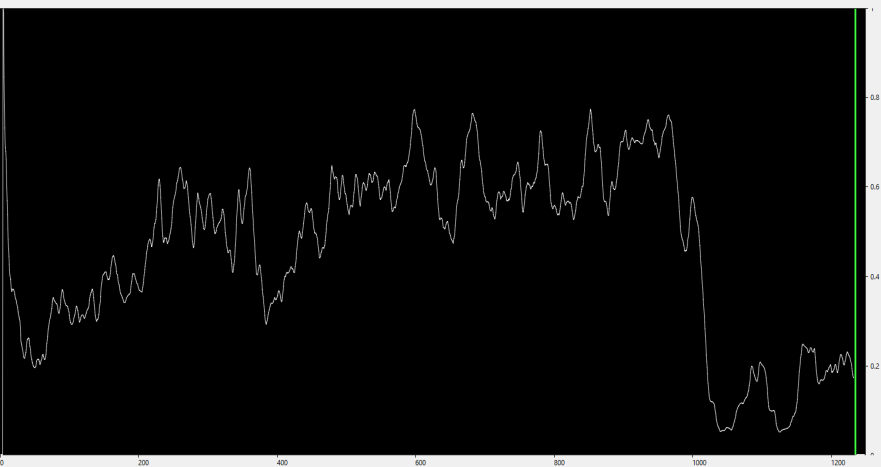

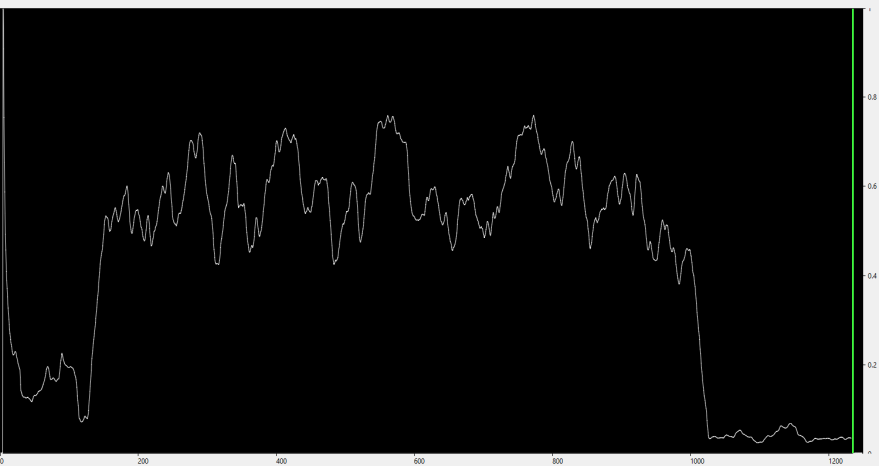
