## Supplementary materials D for "Teaching a computer to assess hypnotic depth: A pilot study"

Patient E had the only frequency band that provided the models with high enough classification accuracy. We assume that the reason for this is that she was the only patient in our study whose experimental conditions were maximally different from session to session. The room and body position were changing: three sessions, including the calibration session, were conducted in a sitting position, and the other three were conducted in a lying position. There were very long intervals between visits (the gap between the first and last sessions was more than 1 year). Most of the experiments with her were accompanied by artifacts in both the low-frequency (approximately 1.5-3 Hz) and the high-frequency (more than 18 Hz) range.

The high-frequency artifacts were due to the myogram, which most likely occurred because of the gradually growing uncomfortable position of the head as a result of increasing relaxation of neck muscles. Because of the artifacts, one of the sessions (#5) with this patient was excluded from the analysis. All of this led to the fact that EEG calibration recording (the first session) gave a classification model that was valid only in the band free of both low and high frequencies, i.e. in the range from 4 to 15 Hz. As seen in Table 3, there was not only high classification accuracy but also very high stability of this accuracy in this band from session to session (SD=1.9). We detected this band at the beginning of our online sessions with her and used it there.

We assume also that the low-frequency artifacts in Patient E could be due to such a common phenomenon as drift (trend) (Driel, 2021). The Predictive curves of her second (on 1.5-45 Hz) and third (on 1.5-45 and 1.5-14 Hz) sessions had a very similar shape to the Native curves, but with the only difference – they were "raised" above the zero value, and it was possible to increase the classification accuracy of these prediction models by subtracting a certain constant value from the feature vector in Scenario-IV (see the notes below Table 3). The lack of classification accuracy in the 1.5-8 and 1.5-14 Hz ranges in Patient C was also associated with low-frequency artifacts.

Consequently, for the BCI paradigm to be used with hypnosis, we could recommend creating conditions with a minimized artifacts probability, i.e. conducting sessions in the same room, body position, etc. We could also recommend pre-training several models based on different frequency bands while calibrating, as was done in our study. If the artifacts are unavoidable, and the initially applied model at the very beginning of the online session demonstrates incorrect waking state prediction, then it is necessary to replace it immediately with another model that is trained on a narrower frequency band, with the exclusion of the slow waves (which may be associated with drift) or high-frequency activity (related to the myogram). Another way to cope with this problem is to correct the scenario by subtracting an empirically selected constant value from the feature vector, thus, eliminating the drift.
